# Machine learning and phylogenetic models identify predictors of genetic variation in Neotropical amphibians

**DOI:** 10.1101/2023.06.15.545105

**Authors:** Luis Amador, Irvin Arroyo-Torres, Lisa N. Barrow

## Abstract

**Aim:** Intraspecific genetic variation is key for adaptation and survival in changing environments and is known to be influenced by many factors, including population size, migration, and life history traits. We investigated genetic variation within Neotropical amphibian species to provide insights into how natural history traits, phylogeny, climatic, and geographic characteristics influence intraspecific diversity.

**Location:** Neotropics.

**Taxon:** Amphibians.

**Methods:** We assembled datasets using open-access databases for natural history traits, genetic sequences, phylogenetic trees, climatic, and geographic data. For each species, we calculated overall nucleotide diversity (π) and tested for isolation by distance (IBD) and isolation by environment (IBE). We then identified predictors of π, IBD, and IBE using Random Forest (RF) regression or RF classification. To incorporate phylogenetic relationships, we fitted phylogenetic generalized linear mixed models (PGLMMs) to predict π, IBD, and IBE.

**Results:** We compiled 4,052 mitochondrial DNA sequences from 256 amphibian species (230 frogs and 26 salamanders), georeferencing 2,477 sequences from 176 species that were not linked to occurrence data. RF regressions and PGLMMs were congruent in identifying range size and precipitation (σ) as the most important predictors of π. RF classification and PGLMMs identified minimum elevation as an important predictor of IBD, and maximum latitude and precipitation (σ) as the best predictors of IBE.

**Main conclusions:** This study unified machine learning and phylogenetic methods and identified predictors of genetic variation in Neotropical amphibians. This approach was valuable to determine which predictors were congruent between methods. We found that species with small ranges or living in zones with less variable precipitation tended to have low genetic diversity. We also showed that Western Mesoamerica, Andes, and Atlantic Forest biogeographic units harbor high diversity across many species that should be prioritized for protection. These results could play a key role in the development of conservation strategies for Neotropical amphibians.

## Introduction

Characterizing genetic variation within species and understanding the processes that maintain that diversity are important goals in evolutionary biology (Ellegren & Galtier, 2016). Genetic variation is essential for natural populations to develop new traits and adapt to future environmental changes (Stange et al., 2021), making it a fundamental aspect of biodiversity conservation efforts (DeWoody et al., 2021). Intraspecific genetic variation includes both the variation within populations and the distribution of spatial genetic variation between populations. These two components provide insights into demographic history and evolutionary processes such as changes in effective population size (N_e_) or the pattern of genetic exchange between populations (Paz-Vinas et al., 2018). Therefore, investigating the forces that drive genetic diversity within species and how this diversity is distributed and influenced spatially, can help explain the processes underlying the patterns of genetic variation that we observe in nature.

Levels of genetic variation vary greatly in natural populations and among species (see Romiguier et al., 2014), however, determinants of intraspecific genetic variation remain poorly understood. For decades, many studies have considered N_e_ to be the most important evolutionary parameter that has a significant impact on genetic variation within and between populations (e.g., Frankham, 1996). For example, small, isolated populations tend to have low genetic variation due to increased genetic drift and reduced gene flow. Under neutral theory, genetic variation is expected to increase with population size (Kimura, 1983), due mainly to reduced genetic drift in large populations (Buffalo, 2021). Several empirical studies have shown results consistent with the neutral theory, in which species with higher population abundances have higher genetic diversity (e.g., Grundler et al. 2019; Hague and Routman, 2016). However, genetic diversity does not necessarily correlate with population size in all cases (e.g., Bazin et al. 2006), and genetic variation levels across species have been observed to be much narrower than their variation in population size (the so-called Lewontin’s paradox; Lewontin, 1974). In nature, genetic variation within species can be influenced by several additional factors, including environmental and intrinsic characteristics (e.g., Nevo, 1978).

Distinct geographic, climatic, and life history factors can determine intraspecific genetic variation and influence the diversification and extinction of populations and species. The range size of a species and the environmental heterogeneity across the species distribution can influence genetic variation. Species with larger ranges are expected to have higher genetic diversity, low inbreeding, and reduced genetic drift because of the direct relationship between geographic range size and population size (see Leffler et al., 2012). For example, studying nine co-distributed amphibian species, García-Rodriguez et al. (2020) found that abiotic factors and geographical features affected the genetic diversity of species in Isthmian Central America. In addition, habitat fragmentation can lead to reduced gene flow and increased genetic differentiation among populations, ultimately reducing genetic diversity within populations because of genetic drift (see Dixo et al., 2009). Life history traits also have an impact on genetic variation, providing a connection between different demographic processes (Duminil et al., 2007). For example, species with shorter lifespans and higher fecundity may have higher genetic diversity because there are more chances for mutation and recombination, and to cover the full gradient of environmental pressures experienced by the species (Romiguier et al. 2014). Body size can be another important predictor of genetic variation within some taxonomic groups. For example, BrünicheDOlsen et al. (2019) found that genetic diversity decreases with increasing body size in Darwin’s finches, with the possible explanation that species abundance is expected to decrease with increasing body size (White et al. 2007). Larger-bodied species may also have higher dispersal ability resulting in differences in spatial genetic variation, as has been demonstrated in bees (López-Uribe et al. 2019) and frogs (Paz et al. 2015).

An improved understanding of these factors can provide insights into how genetic variation is shaped in natural populations. Given the variety of factors and their complex relationships, the approach used to assess potential predictors of genetic diversity is no less important. One approach is to collect new genetic data for a set of taxa of interest and analyze linear models of genetic variation with possible predictors (e.g., Dixo et al., 2009). An alternative approach, which can enable comparisons of many more species, is to use repurposed data that was previously collected for a different research purpose and that can be reanalyzed in a common framework. These two approaches converge in macrogenetics (sensu Blanchet et al., 2017), a field that focuses on the integration of genetic datasets from multiple species at large scales with environmental datasets to identify drivers of intraspecific genetic variation (Leigh et al., 2021). Recently, Pelletier and Carstens (2018) applied a machine learning framework to repurposed georeferenced DNA sequences from more than 8000 species and found that geographic range size and latitude were the most important predictors of genetic structure. Another approach to address how landscapes contribute to the evolution of genetic variation is one that quantifies the effects of geography and ecology using multiple matrix regression with randomization (MMRR; Wang, 2013). Using this approach, Wieringa et al. (2020) examined the impact of geography and ecology on genetic distance of several species in the southeastern United States, finding that geographic or environmental distance were significant factors in most of the species evaluated.

Amphibians have been a preferred study system for several ecological and evolutionary studies because they exhibit a wide variety of natural history traits (e.g., complex life cycles) and distributional patterns (e.g., species with limited dispersal capabilities that lead to high levels of genetic differentiation). Many studies have explored patterns of diversity in amphibians. For example, on a regional scale, a positive latitudinal gradient of taxonomic, functional, and phylogenetic diversity in the Americas has been demonstrated in amphibians (Ochoa-Ochoa et al., 2020). Global genetic diversity patterns also appear to follow a latitudinal gradient in amphibians (Miraldo et al. 2016), but less is known about the predictors of intraspecific variation. In a study of Nearctic amphibian species, the most important predictors of genetic diversity were taxonomic family, number of sequences, and for salamander species (N = 98), those at more northern latitudes had lower genetic diversity (Barrow et al., 2021). In the same region, Schmidt et al. (2022) analyzed microsatellite data from 19 amphibian species; they found that genetic diversity was not predicted by the environmental variables they used and that areas with high species richness also had high genetic structure, but low genetic diversity.

The Neotropical Region includes almost 50% of the world’s total amphibian species, which is greater than any other comparable area on the planet (Menéndez-Guerrero et al., 2020). This exceptionally high diversity is possibly due to a combination of factors, including the geological history of the region (e.g., the formation of the Isthmus of Panama, Andean uplift), climate change, ecological interactions, and biotic diversification (see Elmer et al., 2013; Antonelli, 2022). For decades, the origins and maintenance of diversity in this region has been an important focus of research in amphibian ecology and systematics. For example, Gonzalez-Voyer et al. (2011) found that high-altitude ranges, vascularized ventral skin, and vegetation type correlated significantly with species richness of New World direct-developing frogs. Using glassfrogs as a study model, Castroviejo-Fisher et al. (2014) showed how the climatic and geological history of the Neotropics shaped the diversity of this group, demonstrating that initial uplifts of the Andes and marine incursions are coincidental with divergence times of most glassfrog clades. Recently, Tobar-Suárez et al. (2022) found that amphibian species richness in Neotropical cloud forests increased towards the equator, with frog and caecilian species richness increasing towards lower latitudes, while salamanders showed the opposite pattern. Despite this active research focus on species-level diversity, very little is known about the determinants of intraspecific genetic variation in Neotropical amphibians.

Here, we investigated what factors predict genetic variation within Neotropical frog and salamander species. We gathered genetic sequences, natural history traits, phylogenetic relationships, and climatic and geographic information for each species from open-access databases and literature. We estimated nucleotide diversity (π) and tested for isolation-by-distance (IBD) and isolation-by-environment (IBE) from repurposed DNA sequences. To identify potential hotspots of intraspecific diversity, we built maps of nucleotide diversity across the Neotropics. We then applied machine learning and phylogenetic models to investigate the predictors of π, IBD, and IBE within Neotropical amphibians.

## Materials and Methods

### DNA sequences and associated geographic coordinates

We obtained sequences of the mitochondrial gene cytochrome-b (Cytb) from open access databases. We chose Cytb since this was the most abundant gene in Neotropical amphibian studies and it has informative variation within species (van den Burg et al., 2020; Zeisset and Beebee, 2008). We downloaded sequences from three sources: phylogatR (Pelletier et al., 2022), ACDC (van den Burg et al., 2020) and GenBank (National Center for Biotechnology Information). Although these sources include Cytb sequences for hundreds of species, we chose those species with at least five sequences for further analyses to more adequately represent genetic variation within species (Barrow et al., 2021). Alignments were saved in FASTA format and were edited and aligned using AliView v.1.28 (Larsson, 2014) with the MUSCLE aligner v.3.8.31 (Edgar, 2004) using default settings. Geographic coordinates from each sequence were obtained from phylogatR and GenBank when available, which corresponded to 1,575 sequences (38.87% of the total sequences) from 80 species (31.25% of the total species). We linked 2,477 additional sequences from 176 species to geographic coordinates using either (1) the GeoNames (geonames.org) and GEOLocate (geonames.org) databases by entering the associated location name of the sequence to obtain an approximate occurrence for that sequence; or (2) information from manuscripts (e.g., the original species description, distributional records, or systematic and phylogeographic works; see Appendix S1).

### Metrics of genetic variation

To evaluate genetic diversity within species, we calculated nucleotide diversity (π) from each mtDNA alignment of species and localities using the function nuc.div() in the *pegas* R package (Paradis, 2010). To evaluate spatial genetic variation within amphibian species, we calculated isolation-by-distance (IBD) and isolation-by-environment (IBE) for each species. The ‘raw’ genetic distance (*gendist*; the proportion of sites that differ between each pair of sequences) was calculated using the function dist.dna() in the *ape* R package (Paradis and Schliep, 2019). Topographic distance (*geodist*) was calculated between coordinates associated with each sequence using the function topoDist() implemented in the *topoDistance* R package (Wang, 2020). Environmental distance (*envdist*) was calculated based on 19 bioclimatic variables from the WorldClim database (Hijmans et al., 2005) and tree cover data derived from the ESA WorldCover database (Zanaga et al., 2021). These layers were retrieved at 30-seconds spatial resolution using the function landcover() of the *geodata* R package (Hijmans et al., 2023). We conducted Multiple Matrix Regression with Randomization analyses with the MMRR function in R (Wang, 2013) using the three distances, *gendist, geodist*, and *envdist*, to determine whether each species showed significant IBD and IBE. Regression coefficients of geography (IBD, β*_D_*) and ecology (IBE, β*_E_*) and their significance were calculated after 10,000 permutations. MMRR analyses require datasets with n > 4 and assume that >3 samples show differences in their distances. Based on this premise we used 194 species for this analysis. After classifying each species as having significant IBD or not and significant IBE or not, we used these binary classifications as response variables in subsequent analyses.

### Trait and geographic data compilation

We obtained information on traits from the AmphiBIO dataset (Oliveira et al., 2017), AmphibiaWeb (2022), and specific data from manuscripts (Appendix S1; Table 1). Body size (snout-vent length – SVL), the type of habitat predominantly used by adults, activity (diurnal or nocturnal), and development mode (larval or direct development) were included since these traits are involved in the dispersal capacity of amphibian species and can impact intraspecific genetic differentiation (Hillman et al., 2014). We collected elevational and latitudinal data (mean, maximum, and minimum) from Rolland et al. (2018), who generated this information from the Global Biodiversity Information Facility (GBIF), the International Union for Conservation of Nature (IUCN), and WorldClim (Fick and Hijmans, 2017). From the latter we also obtained temperature (BIO1 = Annual Mean Temperature) and precipitation (BIO12 = Annual Precipitation). We also obtained latitudinal and elevational data from specific manuscripts. Elevational data were also recovered from species accounts in the Amphibian Species of the World 6.1 Online Reference (ASW database; Frost, 2021). The geographical range as a shapefile (.shp) for each species was downloaded from the IUCN portal. We obtained the shapefile for 24 species (e.g., *Allobates hodli*, *Bolitoglossa awajun*) not included in IUCN by calculating minimum convex polygons based on the sequence coordinates and using the function mcp() with the *adehabitatHR* R package (Calenge and Fortmann-Roe, 2023). When necessary, we corrected the polygon for species recently split and not included in IUCN, e.g., *Rhinella marina* – *Rhinella horribilis* (Acevedo et al., 2016). The range area was calculated in square kilometers (km^2^) using the functions shapefile() of the *raster* package and areaPolygon() of the *geosphere* package (Hijmans, 2021; 2022) in R 4.2.2 (R Core Team, 2022). The IUCN Red List Category was also recorded for each species (IUCN, 2022). We tested for multicollinearity of the continuous variables using variance inflation factors (VIF) implemented in the R package *car* (Fox and Weisberg, 2019).

**Table 1.** Variables used in this study to predict nucleotide diversity, isolation-by-distance, and isolation-by-environment in Neotropical amphibians. Variables with an asterisk (*) indicate non-biologically motivated variables and were not used in all analyses. The rationale for considering each variable is provided as an example prediction and is not meant to be an exhaustive explanation.

| Variable | Value | Rationale |
| --- | --- | --- |
| Habitat | Aquatic, Arboreal, Burrowing, Semi-aquatic, Semi-arboreal, Terrestrial | Some habitats may have more natural barriers (e.g., fully aquatic species would have lower dispersal), leading to genetic differentiation of populations. |
| Activity | Diurnal, Nocturnal | Diurnal or nocturnal activity could restrict or promote gene flow, leading to genetic variation between populations. |
| Body size | millimeter (mm) | Larger-bodied species are expected to have higher dispersal ability, facilitating gene flow between populations. Larger-bodied species also may have limits on population abundance. |
| Development mode | Direct, Larval | Species with larval development may have lower dispersal ability since they are tied to water bodies. |
| Sampling effort* | number of sequences >5 | Sampling effort can be considered as a proxy of abundance (population size). Species with more sequences sampled are expected to have higher genetic variation. |
| Sequence length* | number of base pairs | Species with longer CytB fragments may have more variable sites sampled. |
| Elevation (mean, minimum, maximum) | meters above sea level (masl) | Species at higher elevations may experience physical barriers or occur in more isolated populations, increasing genetic differentiation. |
| Latitude (mean, minimum, maximum) | decimal degrees | Latitudinal gradients in climatic variables can influence patterns of distribution and abundance within species. |
| Precipitation<br>(mean, $\sigma$) | precipitation millimeter<br>(mm) | Amphibian diversity is directly affected by precipitation, e.g. species living in areas with high precipitation may have higher abundances. |
| Temperature<br>(mean, $\sigma$) | average temperature ($^{\circ}\text{C}$) | Temperature may not have strong influence on Neotropical species due to the relatively stable and mild temperatures of the region. |
| Range size | area in square kilometers<br>(km<sup>2</sup>) | Species with larger ranges are expected to have higher genetic variation due to larger population sizes and more chances of dispersal. |
| Order | Anura, Caudata | Order-level classification may be associated with characteristics (e.g., reproductive mode, body size) that influence genetic diversity. |
| Family | Anura (17 families),<br>Caudata (Plethodontidae) | Family-level classification may be associated with characteristics (e.g., reproductive mode, distribution) that influence genetic diversity. |
| Biogeographic Unit | 17 units shown in Figure 1a | Biogeographic units are associated with different climates, geologic features, or total area that can affect population sizes and gene flow between populations leading to genetic differentiation. |
| Conservation Status | NE = Not Evaluated,<br>DD = Data Deficient,<br>LC = Least Concern,<br>NT = Near Threatened,<br>VU = Vulnerable,<br>EN = Endangered,<br>CR = Critically Endangered, EX = Extinct | Threatened species with small population sizes are expected to have low genetic diversity; Least concern species are expected to have large range sizes and associated high genetic diversity. |
| Land cover (mean) | 0/1, 1 indicates forest presence | Species living in forested areas (greater number of trees) are expected to have higher abundances and more genetic variation than species living in deforested areas. |

### Geographic framework

To evaluate and visualize how amphibian genetic diversity is distributed in the Neotropical Region, we divided our study area into 17 biogeographic units following the regionalization proposed by Morrone (2014) and with modifications from Antonelli et al. (2018), Josse et al. (2003), and Olson et al. (2001) (Figure 1a). We calculated and mapped intraspecific nucleotide diversity first with a single value of π per species (mean value of π), and second with a value of π per locality within each species. Each locality consisted of at least two individuals, and we assigned sequences to the same locality when they shared the same coordinates. In cases when the sequences were from different geographic coordinates, we combined sequences based on the distance between points (150 km or less) and shared habitat characteristics (e.g., occurring in the same mountain range or river). Although this approach involved arbitrarily choosing a somewhat coarse resolution, it allowed us to maximize the species and geographic area included to provide an initial picture of spatial genetic diversity given the data available. We visualized π for three different resolutions (grid cell size of 150 km^2^, 250 km^2^, and 350 km^2^; Figure 1c, Figure S1b, d) which corresponds to 3640, 1326, and 673 grid cells, respectively. We also visualized the number of sequences per grid cell for the three resolutions to present the sampling effort and distribution of sequences available. All mapping analyses were performed in R. To evaluate the degree of match between the sampling effort (number of observations) and π, we computed the Moran’s I statistic to test for spatial autocorrelation by using the moran.test() function of the *spdep* package (Bivand, 2022).

**Figure 1.**
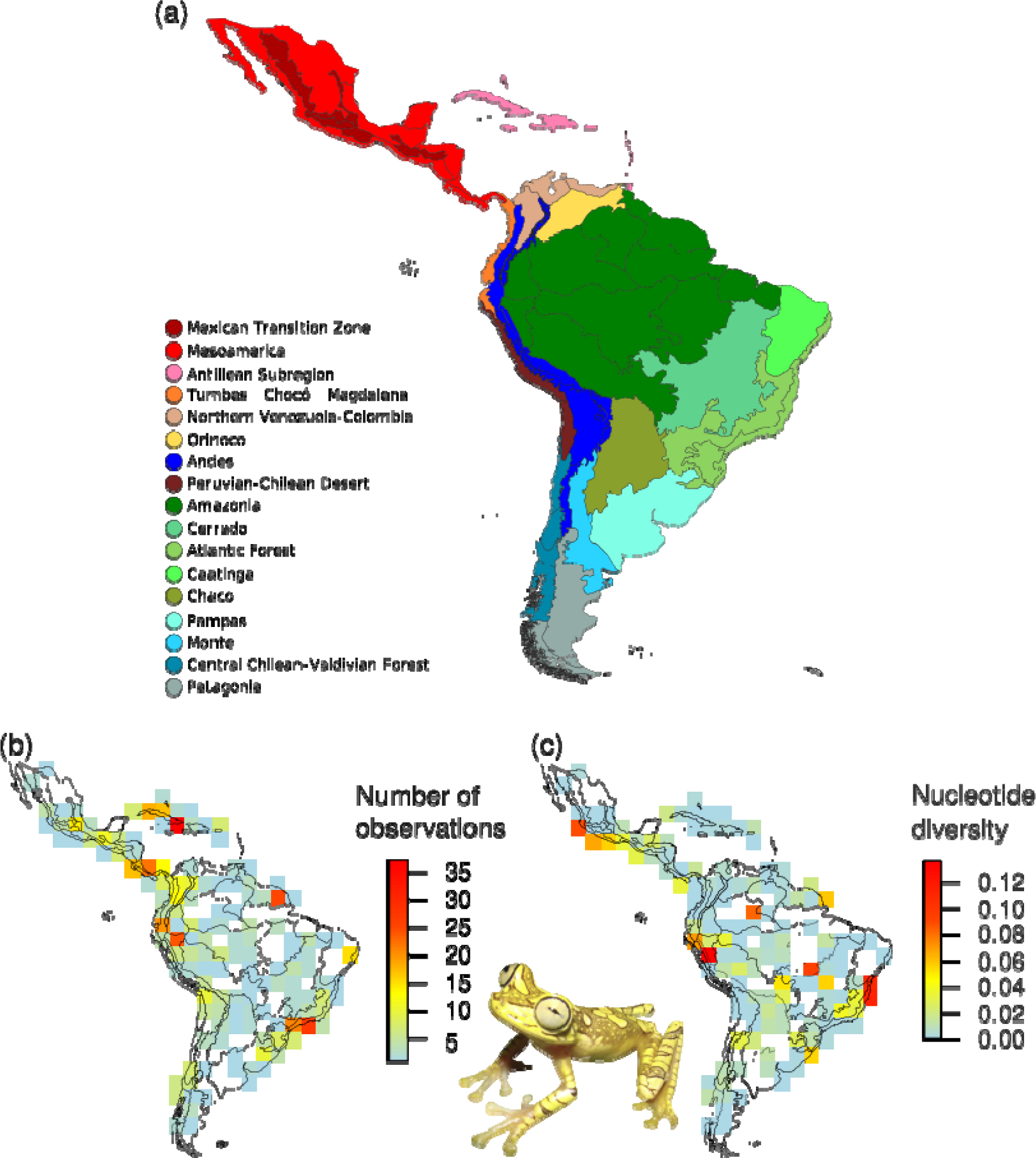
(a) Neotropical biogeographic units were used in this study. Regionalization follows Morrone (2014), with modifications from Olson et al. (2001), Josse et al. (2003), and Antonelli et al. (2018). (b) Number of observations/sequences per grid cell. (c) Amphibian genetic diversity patterns in the Neotropics. The map uses equal-area grid cells of 350 km.

### Identifying predictors of genetic variation with random forests

We used the random forests machine learning algorithm (Breiman, 2001) to build predictive models and identify variables (e.g., body size, habitat, elevation, range size; see Table 1) that are important predictors of π, IBD, and IBE in Neotropical amphibians. RF uses independent variables to create many individual decision trees (a forest) that act as an ensemble to predict a response. Each decision tree in the random forest uses a subset of the independent variables and returns a response, and variable importance is determined based on the increase in model error when that variable is not included (Kabacoff, 2015). RF regression (for π) and classification (for IBD and IBE) models were created using the randomForest() function in the *randomForest* R package (Liaw and Wiener, 2002), with 5000 trees and 100 permutations. We split the data into training (90%) and test (10%) datasets and created RF models with the training data. We used the tuneRF() function to find the optimal mtry value (number of variables to randomly sample as candidates at each split), and a new model was built using the best mtry value. We made predictions on the training and test (unseen data values in the models) datasets that were evaluated with the Mean Squared Error (MSE) and R-squared (R^2^) metrics in the RF regression analysis and with the confusion matrix in RF classification analyses. Relative importance for each independent variable was measured and printed using the importance() function in the *randomForest* package, and visualized with the vip() function of the *vip* R package (Greenwell and Boehmke, 2020) for each model. To evaluate the effect the number of sequences per species and the number of base pairs (bp) in each alignment might have on the findings, we also created RF models with two different reduced datasets: (1) a dataset with at least 10 sequences per species (114 species), and (2) a dataset with at least 400 bp in each species alignment (197 species).

### Testing predictors of genetic variation with phylogenetic comparative methods

Random forests can handle large numbers of variables to build predictive models, but do not explicitly incorporate phylogenetic relationships (other than taxonomic level as a possible predictor). Therefore, we also fitted phylogenetic generalized linear mixed models (PGLMMs) to determine relationships between natural history traits, geographic and climatic variables, and genetic variation of Neotropical amphibians. We pruned the phylogenetic tree from Jetz and Pyron (2018) to create a phylogeny that includes only the species in our dataset using the phylo4() function in the *phylobase* R package (Hackathon et al., 2020). This resource is the most complete phylogeny available (7,238 species), covering ≈83% of the known amphibian diversity. Of the 256 species in our dataset, 15 species were not included in the phylogeny of Jetz and Pyron (Appendix S2), therefore for phylogenetic analyses we used a subset consisting of 241 species. We mapped π as a continuous trait using the contMap() function in the *phytools* R package (Revell, 2012), and the distribution of IBD and IBE were mapped at the tips on the tree. We tested for phylogenetic signal in π using two metrics, Pagel’s λ (lambda; Pagel, 1999) and Blomberg’s K (Blomberg et al., 2003), calculated with the phylosig() function implemented in *phytools*. We tested for phylogenetic signal in IBD and IBE using the phylo.d() function that calculates the D statistic, a measure of phylogenetic signal in a binary trait (Fritz and Purvis, 2010), implemented in the R package *caper* (Orme et al., 2018).

Finally, we used PGLMMs (Ives and Helmus, 2011) implemented in the *MCMCglmm* R package (Hadfield, 2010) to investigate relationships between a subset of the predictors and three responses: (1) π, (2) IBD, and (3) IBE, while accounting for phylogeny. For these models, we included species ‘random effects’, which account for the variability caused by species-specific effects, and phylogenetic random effects, which consider the phylogenetic relationship between species, by transforming the phylogeny into a variance-covariance matrix of relatedness between species (Garamszegi, 2014). For each response variable (π, IBD, and IBE), we compared 10 models that were generated with different combinations of predictors based on the results of RF analyses (Table S2). These models included: (1) an intercept-only null model, (2) a model with both species and phylogenetic random effects and no predictors, (3) a model with six predictors and no random effect, (4) a model with six predictors and species as a random effect, (5) a model with six predictors and a phylogenetic random effect, (6) a model with six predictors and species and phylogenetic random effects, (7) a model with 12 predictors and no random effects, (8) a model with 12 predictors and a species random effect, (9) a model with 12 predictors and a phylogenetic random effect, and (10) a model with 12 predictors and with both species and phylogenetic random effects. For Bayesian model selection, we used the deviance information criterion (DIC).

## Results

### Data compilation

We compiled 4,052 Cytb mtDNA sequences from 256 Neotropical amphibians (Figure S2), of which only 80 species had associated occurrences (we retrieved occurrences for 50 species from phylogatR and 30 species from GenBank). Occurrences for the sequences of the other 176 species were recovered by us in the present work by retrieving coordinates from published literature or by georeferencing localities. To visualize sampling effort across the region, we mapped the number of sequences per grid cell (Figure 1b, Figure S1a, c). There was weak positive spatial autocorrelation of number of sequences (Moran’s I = 0.0569, P-value = 0.019), this suggest that similar number of sequences values are somewhat clustered together but is not a strong pattern in the region. The total occurrences represented all 17 Neotropical biogeographic units (Figure S3). The final dataset included 230 frogs in 17 families and 26 salamanders in family Plethodontidae, three response variables (π, IBD, and IBE), and 22 predictor variables (Table 1, Table S1). We did not detect correlation among predictor variables in the model (VIF values < 2) except for the three elevation variables (mean, minimum, and maximum) (Figure S4).

### Distribution of genetic variation in the Neotropics

The calculated π ranged from 0 (10 species) to 0.156 (Boquete rocket frog, *Silverstoneia nubicola*, family Dendrobatidae) with a mean of π = 0.025 (Table S1). When we visualized nucleotide diversity per locality, we found higher π values in western Mesoamerica, central Andes, Atlantic Forest, and a few areas of the Amazon region, compared to their adjacent biogeographic units. On the other hand, the southern Andes and the southern portion of South America showed low genetic diversity (Figure 1c). Mean values of π per species were higher in southern Mesoamerica, the Chocó region, northern Andes, and Atlantic Forest region (Figure S5). Spatial autocorrelation analysis suggests that is likely that π values are randomly distributed in the geographic space (Moran’s I = 0.0662, P-value = 0.009). With respect to spatial genetic variation, we found that 89 Neotropical amphibian species showed significant IBD, while 105 species did not (Table S1). Species following an IBD pattern were mostly found in Mesoamerica (n = 18), the Antillean Subregion (n = 13), the Atlantic Forest (n = 13), and the Andes region (n = 13). The Amazon included the most species that did not present IBD (n = 28), followed by the Atlantic Forest and Andes region (Figure S6). We found that most species did not show IBE (n = 124) and only 70 species showed significant IBE. Species with IBE were found mostly in the Antillean region (n = 14), Andes (n = 13), and Amazonia (n = 12). The Amazon region also included the most species with no IBE patterns (n = 27), followed by the Atlantic Forest and Mesoamerica (Figure S7)

### Important predictors of genetic variation based on random forests

Random forest regressions showed that range size was the most important predictor of π for Neotropical amphibians (Figure 2a). Precipitation standard deviation (σ) was another important predictor of π, followed by body size and mean temperature (Figure 2a). The overall variance explained by the RF regression model shown was 20.31%. Random forest classification found that body size was the most important predictor of IBD, followed by mean precipitation, IUCN rank, and minimum elevation (Figure 2b). The most important predictor for IBE was maximum latitude, followed by precipitation (σ) and latitude (mean and minimum) (Figure 2c). The model error for the RF classification model for IBD was 37.21% and for IBE was 40.21%. When we evaluated the effect of the sampling effort (number of sequences) per species (at least 10 sequences), and the sequence length (number of base pairs, bp) in each alignment (at least 400 bp) on our results, we found that range size was the best predictor of π for all models analyzed (Figure S8) as in the model using the complete dataset. For IBD, body size remained one of the most important predictors for the ‘at least 10 sequences’ dataset, but not the ‘at least 400 bp’ dataset. Instead, mean temperature was the most important predictor of IBD in all the models using reduced datasets (Figure S9). Mean temperature was also the best predictor of IBE in the ‘at least 400 bp’ dataset, while microhabitat was the best predictor for the ‘at least 10 sequences’ dataset (Figure S10). For IBE, latitude was still among the most important predictors for both reduced datasets.

**Figure 2.**
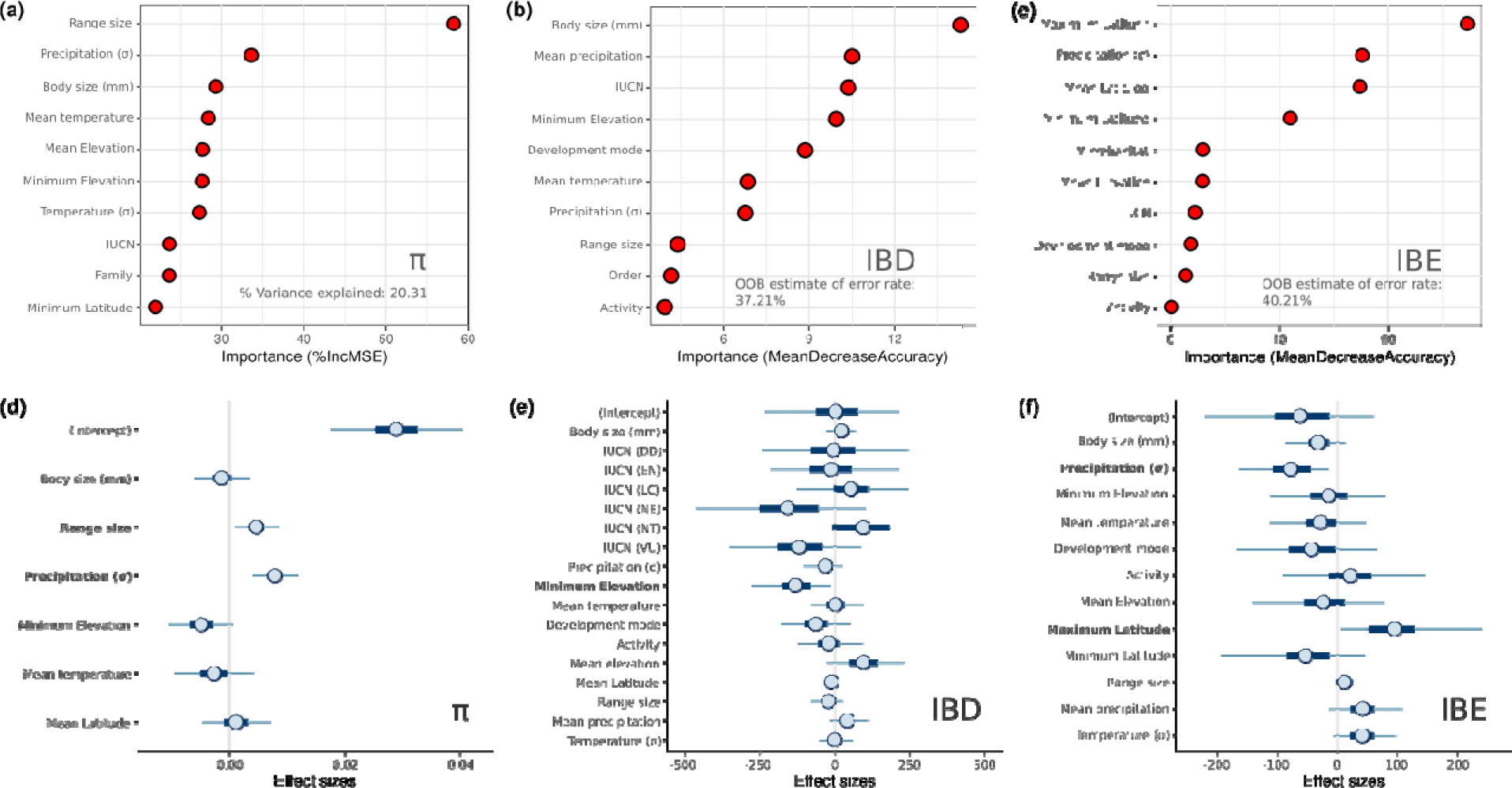
Top ranked predictors according to their variable importance for random forest models in explaining (a) π, (b) IBD, and (c) IBE. Effect sizes of π (d), IBD (e), and IBE (f) across predictors in *MCMCglmm* analyses. The associated 95% credible intervals (CIs) do not cross zero for range size and precipitation in π, for minimum elevation in IBD, and precipitation and maximum latitude in IBE; indicating that these predictors are statistically significant.

### Relationships between predictors and genetic variation based on PGLMMs

Phylogenetic generalized linear mixed models using *MCMCglmm* showed that the best fit model for π was the one with a reduced number of predictors and with both species and phylogenetic random effects (Table 2). This model indicated that range size and precipitation (σ) predicted π, consistent with our random forest results. Both range size and precipitation (σ) had positive relationships with π (Figure 2d). Neotropical amphibian species with larger ranges and living in areas with more variable precipitation tended to have higher π (Figure 3a-c). For IBD, the best-fit *MCMCglmm* model was the full model with 12 predictors and species and phylogenetic random effects (Table 2). *MCMCglmm* also identified minimum elevation as a significant predictor of IBD, and their relationship was negative (Figure 2e). Neotropical amphibians living at higher elevations tended to have no IBD, and those with significant IBD tended to live at lower elevations (Figure 3d). Similar to RF analyses, precipitation (σ) and maximum Latitude predicted IBE in PGLMM analyses (Figure 2f, Table 2). Species living at southern latitudes in the Neotropics tended to have no IBE, contrary to amphibians in northern latitudes that tended to exhibit IBE (Figure 3e). Species following an IBE pattern mostly occur in areas with lower precipitation (σ) (Figure 3f). *MCMCglmm* results were verified by checking model diagnostics using trace and density plots of the MCMC samples.

**Figure 3.**
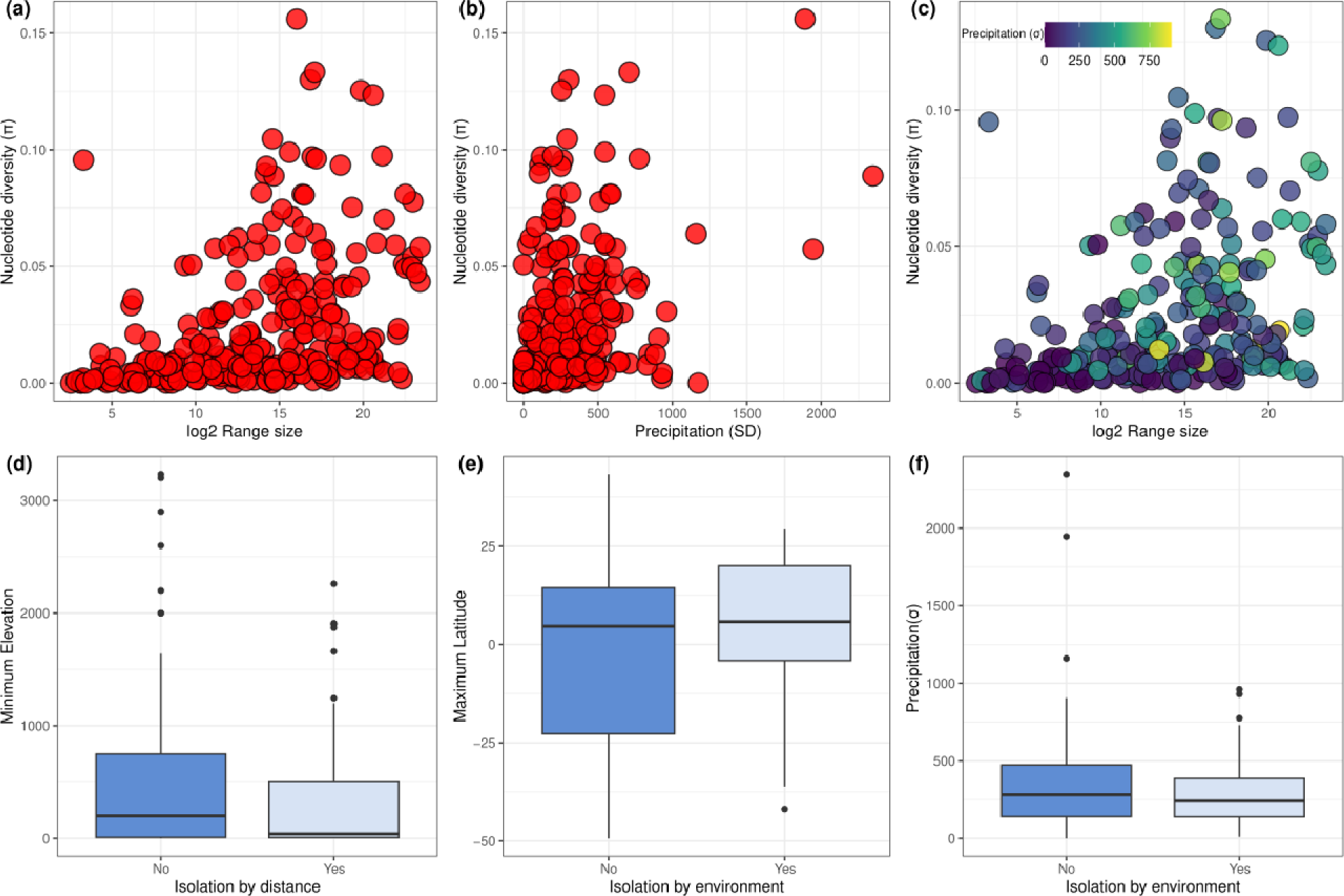
(a) Range size (log2) and mtDNA π for all amphibian species in our dataset; (b) precipitation (σ) and π relationship; (c) Range size (log2) and π relationship without outliers, the colors indicate precipitation (σ) of occurrences in each species. (d) Minimum elevation for species with and without of isolation-by-distance (IBD). (e and f) Maximum latitude and precipitation (σ) for species with and without isolation-by-environment (IBE). Each box-whisker plot indicates the median (bold lines), the interquartile range (boxes), and dots represent outliers.

**Table 2.**
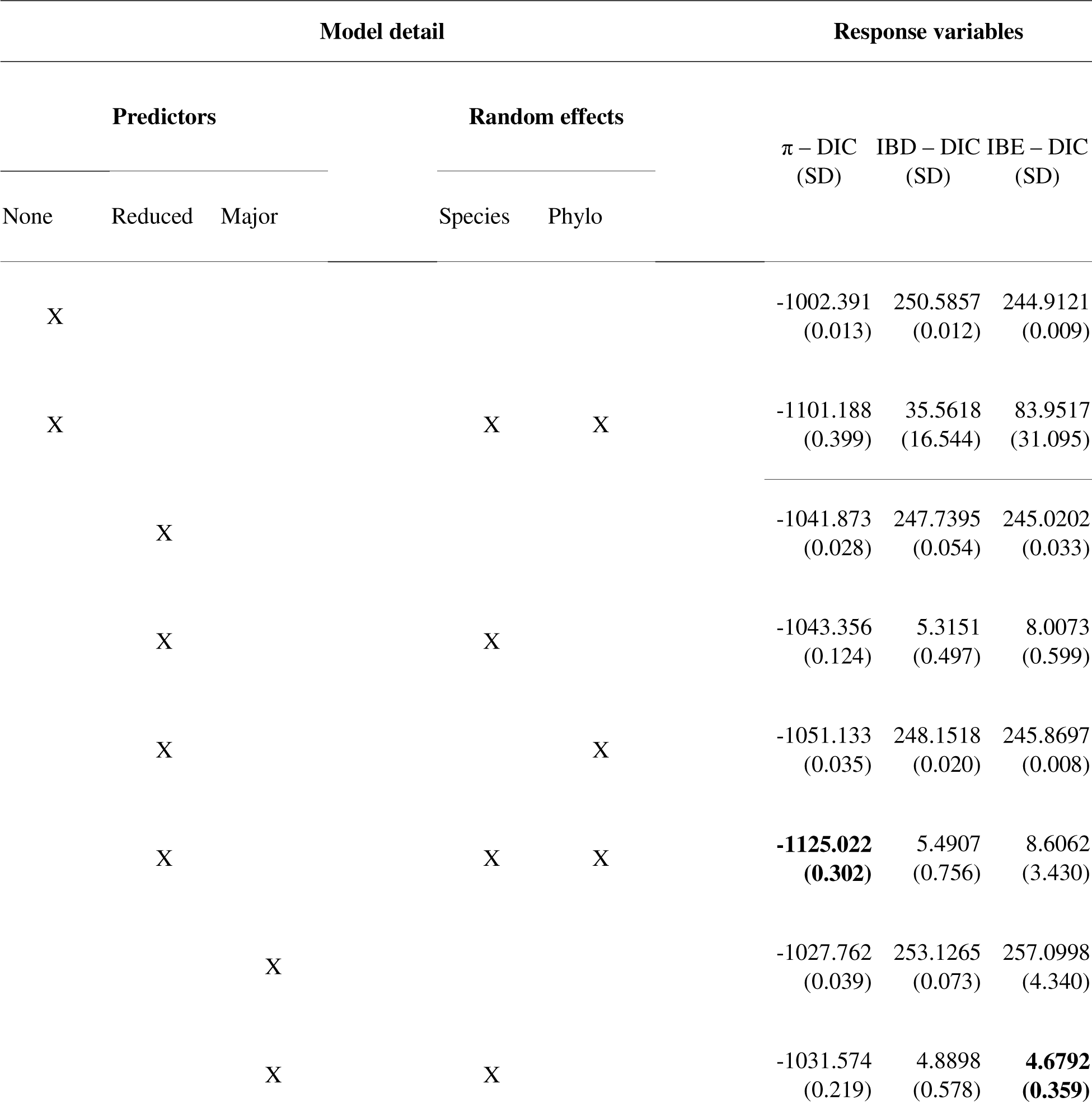
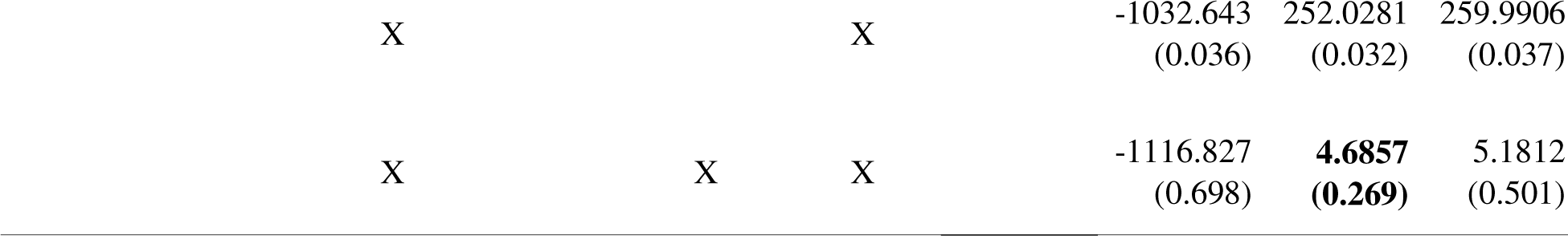
Model comparison with deviance information criterion (DIC) scores from each MCMCglmm model for nucleotide diversity (π), isolation-by-distance (IBD), and isolation-by-environment (IBE). Comparisons were made with a major and reduced number of predictors and without predictors; with no random effects and with species and phylogenetic (phylo) random effects. For each model, four different runs were performed; we present the average DIC score with the standard deviation (SD).

### Testing phylogenetic signal

We found significant phylogenetic signal in π (Pagel’s λ = 0.7684, P-value (based on LR test) < 0.0001; Blomberg’s K = 0.1509, P-value (based on 1000 randomizations) = 0.002) (Figure 4; Figure S11a, b). We found clusters of closely related genera that had dissimilar π values; for example, within Bufonidae, *Atelopus* species had low π (average = 0.006, range: 0.001-0.010) while toads of genus *Rhinella* presented a higher average π (average = 0.020, range: 0.006-0.054). The same pattern was also observed in poison frogs of the family Dendrobatidae, where *Ameerega* species showed high π (average π = 0.024; range: 0.003-0.059) in contrast to species in the genus *Andinobates* which had very low π (average π = 0.003; range: 0.000-0.009). We did not find significant phylogenetic signal in IBD or IBE, with values of D that were greater than 1 and were overdispersed compared to a Brownian threshold model (Estimated D_IBD_ = 1.008, P-value = 0.53; Estimated D_IBE_ = 1.047, P-value = 0.685) (Figure S11c, d).

**Figure 4.**
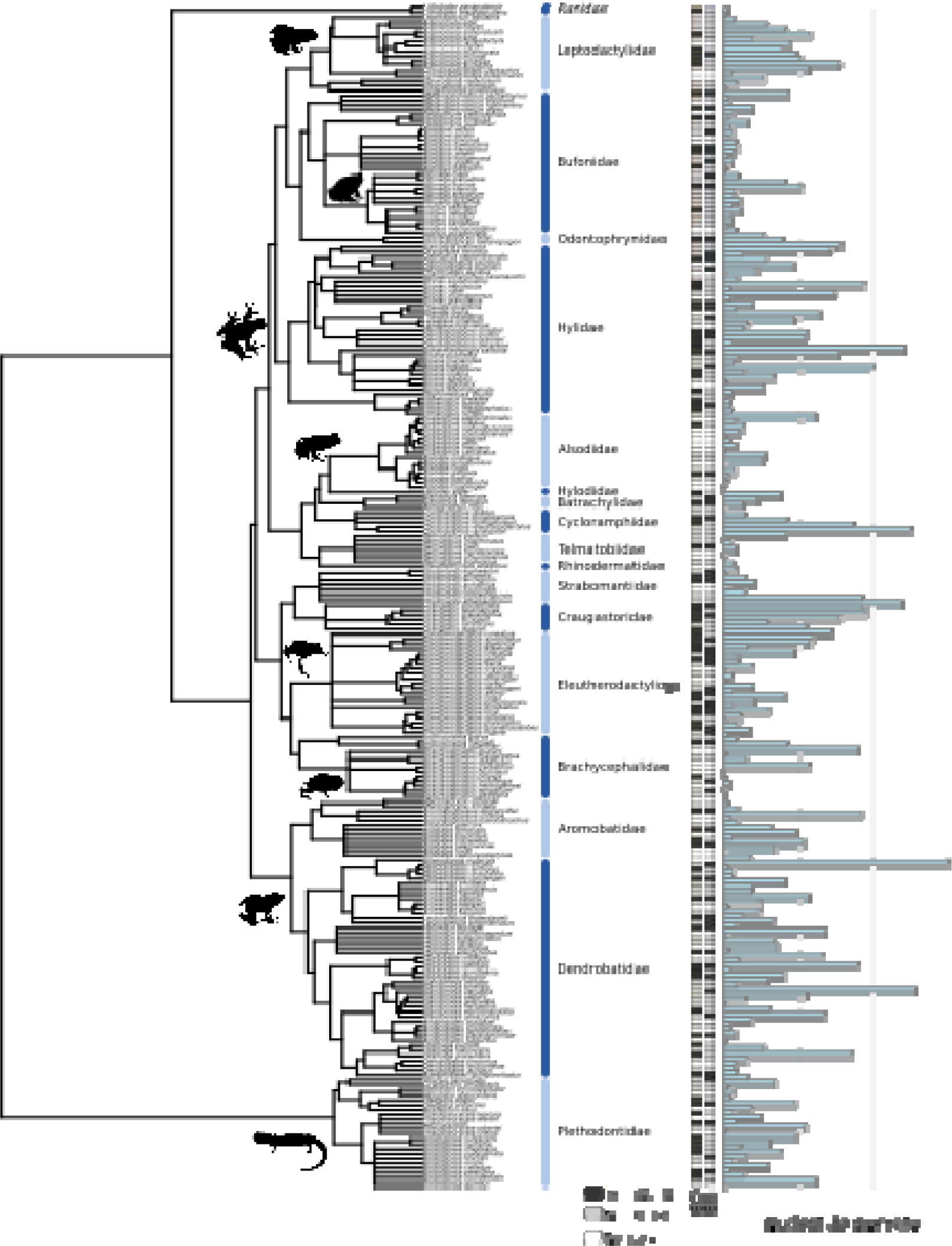
Phylogeny of Neotropical amphibians (downloaded and subset from Jetz and Pyron, 2018) with values of CytB π presented as light blue bars, and presence (black dots) or absence (grey dots) of IBD and IBE in Neotropical amphibians. Silhouettes of frogs and salamanders were obtained from phylopic.org.

## Discussion

In this study we investigated the predictors of genetic variation (π, IBD, and IBE) in Neotropical amphibians using repurposed data including mtDNA sequences, natural history traits, and geographic information gathered from open-access databases. Our analyses revealed that geographic range size, precipitation, elevation, and latitude were significant predictors of different aspects of genetic variation within species. Specifically, we found that amphibian species inhabiting smaller ranges and places with lower variation in precipitation had lower intraspecific π (Figure 3a, b); species living at higher elevations (e.g., mountain ranges) tended not to exhibit IBD; and species living in southern latitudes tended not to exhibit IBE (Figure 3d, e).

Geographic range size was the top predictor of π, but did not predict IBD and IBE, i.e., Neotropical amphibian species with and without IBD or IBE can inhabit small or large geographic ranges. We suspect that environmental heterogeneity within the geographic range rather than range size may have a greater influence on IBD and IBE patterns of species in the Neotropics, as has been observed in other vertebrate studies (e.g., Quiroga-Carmona and D’Elía, 2022). Geographic range size has also emerged as an important predictor of genetic variation in other taxonomic groups such as squamates (Larkin et al., 2023) and Darwin’s finches (Brüniche-Olsen et al., 2019). Interestingly, the relationship between genetic diversity and geographic range is not always straightforward, as evidenced by the case of butterflies (Mackintosh et al., 2019) and other non-model animal species (Romiguier et al., 2014). These discrepancies highlight the need for continued study of a variety of taxonomic groups and regions to better understand the ecological and geographic context of genetic variation. Geographically restricted species are expected to have less genetic variation, which may also indicate an increased risk of extinction (Levy et al., 2016). In amphibians, Caviedes-Solis et al. (2020) found that Neotropical treefrogs living in high elevations were more likely to be classified with threatened status. Our analysis of Neotropical amphibians from several taxonomic families confirmed expected differences between geographic range sizes based on IUCN conservation status, with Least Concern (LC) amphibian species occurring in larger ranges and Data Deficient (DD), Endangered (EN) and Critically Endangered (CR) species occurring in smaller ranges (Figure S12). Based on our estimates, however, IUCN conservation status did not have as clear of an association with π (Figure 2a, Figure S12), suggesting that IUCN status may not currently capture this important parameter for population persistence. Regardless, populations of threatened species with low π such as harlequin frogs of the genus *Atelopus* should be closely monitored.

Precipitation and temperature, along with associated gradients of elevation, has significant impacts on genetic variation across animal species (De Kort et al., 2021). Our study identified minimum elevation as a significant predictor of IBD (and it was also one important predictor of π), and precipitation as one of the best predictors of <u>π,</u> IBD, and IBE (Figure 2). We observed that species inhabiting higher elevations did not usually exhibit IBD (see Figure 3d), while those living in lower elevations tended to have both significant IBD patterns and larger geographic range sizes. The absence of IBD patterns, and in turn low π, in species living at higher elevations could be attributed to their smaller ranges, leading to smaller population sizes. Our findings partially agree with those reported by Pelletier and Carstens (2018), who found that geographic range size, elevation, and latitude predicted IBD in several taxonomic groups. In our case, latitude (maximum) was the most important predictor of IBE. Species with a significant IBE pattern occurred at higher latitudes than species without IBE. Although latitude was not a significant predictor of π, analyzing mean latitude with π and geographical range size revealed that amphibians near the equator have larger ranges, and diversity increases near the equator and slowly decreases towards higher latitudes (Figure S13). This pattern of genetic diversity forms a plateau around the equator, a pattern noted by Pereira (2016) in his perspective about a global study mapping amphibian genetic diversity by Miraldo et al. (2016).

The nucleotide diversity map of Neotropical amphibians clearly shows areas of high π, such as Chocó, northern Andes, and Atlantic Forests in South America, and an area of high diversity in the Mesoamerica region located in Central America. Biogeographic units identified with higher π values are areas that also have high precipitation rates. Therefore, precipitation could explain genetic variation in the Neotropics; however, this hypothesis needs to be tested in future studies with more species homogeneously distributed throughout the region. Other studies have previously shown the important role of annual precipitation in explaining species richness, phylogenetic diversity, and functional diversity in Neotropical amphibians (Amador et al., 2019; Ochoa-Ochoa et al., 2019). Differences in research and collecting efforts throughout the Neotropical region could be complicating the interpretation of our genetic diversity map, mainly because we do not have an equilibrated sampling and our repurposed dataset has a high concentration of occurrence points in certain regions (e.g., Amazonia, Atlantic forests or Mesoamerica). In addition, we were not able to recover more than five CytB sequences (our minimum number of sequences per species) for any Gymnophiona species, nor were we able to recover genetic information for centrolenid frogs, one of the most diverse taxa in the Neotropics. We were also unable to recover target sequences for marsupial frogs of the family Hemiphractidae or for Neotropical microhylid frogs. These two groups have high species richness but are lacking in available evolutionary studies. To alleviate some of these sampling issues, future studies should consider comparisons of other mitochondrial and nuclear genes and ideally standardize the markers sequenced across multiple species.

Comparing our results with those for Nearctic amphibians (Barrow et al. 2021), no differences in average π within species were evident (Nearctic amphibians: average π = 0.028, n = 299; Neotropical amphibians: average π = 0.025, n = 256). However, the highest values of intraspecific π were found in several Neotropical species; for example, only three Nearctic species had π values > 0.09 compared to 13 Neotropical species with similar or higher values. This disparity between Neotropical and Nearctic species could relate to differences in demographic history between regions or could be explained by bias in taxonomic practices (see Chek et al., 2003). It is possible that the higher genetic variation we observed in Neotropical amphibians is partially due to taxonomic under-splitting, which could lead to severe conservation implications for this group. We found very high values of π (e.g., > 0.09) for several species, suggesting the possibility of cryptic species in our dataset. At least nine of the species with high π values (Table S3) have been considered as species complexes or cryptic species in previous taxonomic studies (e.g., *Pristimantis altamazonicus* – Ortega-Andrade et al., 2017; *Anomaloglossus degranvillei* – Vacher et al., 2017; *Phyllobates lugubris* – Márquez et al., 2020; *Dendropsophus decipiens* – Oliveira et al., 2021). Higher environmental and climatic heterogeneity may lead to diversification dynamics with higher speciation and lower extinction rates supporting rapid evolution in the Neotropics (Brown, 2014). Under this scenario, species would have accumulated faster towards the present mainly due to recent geological and climatic perturbations (e.g., the elevation of the Andes) (Meseguer et al., 2022). The potential under-split species in our data set are therefore expected to be young or recently diverged, which is one of the mechanisms causing cryptic diversity (Fišer et al., 2018). We interpret the results of genetic variation of these species with caution because they may represent multiple taxa, for example, if we were able to split cryptic species in separate species, range sizes would be smaller impacting the levels of intraspecific genetic diversity.

We found that levels of π varied among families, with species in certain families such as Dendrobatidae and Hylidae having the highest π estimates and Ranidae, Rhinodermatidae, and Telmatobiidae having the lowest (Figure S14). In contrast, the presence of spatial genetic variation (IBD and IBE) within species appears to be more randomly distributed throughout the phylogeny of Neotropical amphibians. These findings highlight the value of employing different methodological frameworks as we did in this study. With the growing size and complexity of biodiversity datasets, machine learning methods such as random forests prove valuable in identifying potential predictors, even using both complete and reduced datasets. Combining these methods with phylogenetically informed models allowed us to gain a deeper understanding of the relationships between predictors and genetic variation within species. For example, our RF (regression and classification) and *MCMCglmm* models were consistent in identifying range size and precipitation as the best predictors of genetic variation (Table 2; Figure 2).

Our study provides valuable insights into the distribution of genetic variation in Neotropical amphibians and identifies important predictors of intraspecific genetic variation. These findings underscore the importance of considering both nucleotide diversity and spatial genetic variation in the conservation and management of Neotropical amphibian populations. For example, the results demonstrate the importance of preserving forest areas, especially in biogeographic areas where the intraspecific π is very low. This information is also valuable to assess the conservation status of Neotropical amphibian species and consider the impact of threats these taxa could be facing in the future. The current distribution of genetic variation could play a key role in the development of targeted conservation strategies for amphibian species, particularly considering the diverse life histories observed among Neotropical amphibians. For example, certain species within the genera *Atelopus* or *Telmatobius*, which are highly endangered groups, require specific conservation measures focused on preserving aquatic habitats (e.g., streams, ponds, or lakes). These habitats serve as crucial breeding grounds where frogs lay their eggs in shallow water, making their conservation of paramount importance. To address these and other subjects inherent to amphibian ecology and evolution, we suggest that future work should include more information such as genome-scale data and where possible, add more species within the region and globally.

## Acknowledgements

This work was supported by a grant to Lisa Barrow from the U.S. National Science Foundation (DEB-2112946). Thank you to the associate editor and two reviewers for their suggestions that helped to improve this manuscript. We thank the members of the Amphibian & Reptile Biodiversity Lab (The University of New Mexico) for their valuable comments on earlier versions of this manuscript, Bryan Carstens (The Ohio State University) for early access to the phylogatR database, and Jorge “Colo” Ortega for use of the photo of *Boana picturata* in Figure 1.

## Data Availability

Supplementary table S1 and additional supplementary files (e.g., appendices, fasta sequences used for the analyses) are available on Dryad. https://datadryad.org/stash/share/mAYGsdKODg3I8ouXADLggPVcTkVkU7Nqa1wGPl80jiQ

## Biosketch

Luis Amador is an evolutionary biologist, currently studying the determinants of global amphibian genetic/genomic diversity as a postdoctoral fellow in the Amphibian and Reptile Biodiversity Lab (ARBL) at the University of New Mexico (UNM). Irvin Arroyo-Torres contributed to this work as an undergraduate researcher and is pursuing graduate studies in ecology and conservation at UNM. Lisa Barrow is Curator of Amphibians and Reptiles at the Museum of Southwestern Biology and principal investigator of ARBL at UNM, which focuses on research in evolution, ecology, and conservation of herpetofauna.

## Author contributions

L.A. and L.N.B. conceived the ideas; L.A. and I.A.T. assembled the data; L.A. analyzed the data; and L.A. and L.N.B. led the writing.

## Appendices

Appendix S1 Literature used to retrieve occurrences associated with DNA sequences and geographical data.

Appendix S2 Species not included in the phylogenetic analyses. Appendix S3 Supplementary tables.

Appendix S4 Supplementary figures.

