## Appendix for "Machine learning and phylogenetic models identify predictors of genetic variation in Neotropical amphibians" (from another version of this preprint: version 1, 2023-06-15)

**Journal of Biogeography**

### Supporting information

**Appendix S1** Materials and methods – supplementary information

### Supplementary tables

**Table S1.1.** The data set used in this study. Nucleotide diversity ($\pi$) and Isolation-by-Distance (IBD) were used as dependent variables in the analyses. Lar=Larvae, Dir=Direct, DD= Data Deficient, LC=Least Concern, NT=Near Threatened, VU= Vulnerable, EN=Endangered, CR= Critically Endangered, EX= Extinct, NE= Not Evaluated. * This table can be found in the figshare online repository https://doi.org/10.6084/m9.figshare.23520384.v1

### Literature used to retrieve occurrences associated with DNA sequences and geographical data.

Anderson, M. T., & Mathis, A. (1999). Diets of Two Sympatric Neotropical Salamanders, *Bolitoglossa mexicana* and *B. rufescens*, with Notes on Reproduction for *B. rufescens*. Journal of Herpetology, 33(4), 601. https://doi.org/10.2307/1565576

Angulo, A., Cocroft, R. B., & Reichle, S. (2003). Species identity in the genus *Adenomera* (Anura: Leptodactylidae) in Southeastern Peru. Herpetologica, 59(4), 490–504. https://doi.org/10.1655/20-104

Angulo, A., & Icochea, J. (2010). Cryptic species complexes, widespread species and conservation: Lessons from Amazonian frogs of the *Leptodactylus marmoratus* group (Anura: Leptodactylidae). Systematics and Biodiversity, 8(3), 357–370. https://doi.org/10.1080/14772000.2010.507264

Baldo, D., Borteiro, C., Kolenc, F., Rosset, S., Prigioni, C., & Martínez Debat, C. (2012). The taxonomic status of *Melanophryniscus orejasmirandai* Prigioni & Langone, 1987 “1986” (Anura: Bufonidae). Zootaxa, 3235(1), 1.

Barrasso, D. A., & Basso, N. G. (2019). Low genetic divergence but many names in the endemic Patagonian frogs of the genus *Atelognathus* (Anura, Batrachylidae): A molecular genetics and morphological perspective. Journal of Zoological Systematics and Evolutionary Research, 57(2), 383–399. https://doi.org/10.1111/jzs.12259

Barrasso, D. A., Úbeda, C. A., Cotichelli, L., & Basso, N. G. (2022). On the presence of *Alsodes coppingeri* (Anura, Alsodidae) in Argentina, with comments on other southern *Alsodes*. Neotropical Biodiversity, 8(1), 21–30. https://doi.org/10.1080/23766808.2022.2029321

Blotto, B. L., Nuñez, J. J., Basso, N. G., Úbeda, C. A., Wheeler, W. C., & Faivovich, J. (2013). Phylogenetic relationships of a Patagonian frog radiation, the *Alsodes* + *Eupsophus* clade (Anura: Alsodidae), with comments on the supposed paraphyly of *Eupsophus*. Cladistics, 29(2), 113–131. https://doi.org/10.1111/j.1096-0031.2012.00417.x

Borteiro, C., Gobel, N., Kolenc, F., Laufer, G., Martínez Debat, C., & Ubilla, M. (2018). Skin-mates or neighbors? A seasonal study of amphibian chytrid and dermocystid infection in *Boana pulchella* (Anura: Hylidae). Cuadernos de Herpetología, 32(2). https://doi.org/10.31017/CdH.2018.(2018-012)

Brown, J. L., Twomey, E., Amézquita, A., De Souza, M. B., Caldwell, J. P., Lötters, S., Von May, R., Melo-Sampaio, P. R., Mejía-Vargas, D., Perez-Peña, P., Pepper, M., Poelman, E. H., Sanchez-Rodriguez, M., & Summers, K. (2011). A taxonomic revision of the Neotropical poison frog genus *Ranitomeya* (Amphibia: Dendrobatidae). Zootaxa, 3083(1), 1. https://doi.org/10.11646/zootaxa.3083.1.1

Charrier, A., Correa, C., Castro, C., & Méndez, M. A. (2015). A new species of *Alsodes* (Anura: Alsodidae) from Altos de Cantillana, central Chile. Zootaxa, 3915(4), 540. https://doi.org/10.11646/zootaxa.3915.4.5

Correa, C., & Durán, F. (2019). Taxonomy, systematics and geographic distribution of ground frogs (Alsodidae, *Eupsophus*): A comprehensive synthesis of the last six decades of research. ZooKeys, 863, 107–152. https://doi.org/10.3897/zookeys.863.35484

Correa, C., Morales, J., Schussler, C., & Ortiz, J. C. (2020). An enigmatic population of *Alsodes* (Anura, Alsodidae) from the Andes of central Chile with three species-level mitochondrial lineages. Mitochondrial DNA Part A, 31(1), 25–34. https://doi.org/10.1080/24701394.2019.1704744

Correa, C., Zepeda, P., Lagos, N., Salinas, H., Palma, R. E., & Vásquez, D. (2018). New populations of two threatened species of *Alsodes* (Anura, Alsodidae) reveal the scarce biogeographic knowledge of the genus in the Andes of central Chile. Zoosystematics and Evolution, 94(2), 349–359. https://doi.org/10.3897/zse.94.25189

Cuevas, C. C., & Sanhueza, R. (2020). Geographic boundaries and natural history notes of the microendemic endangered frog *Eupsophus migueli* Formas, 1977 (Alsodidae) in the Mahuidanche Range, southern Chile. ZooKeys, 929, 79–92. https://doi.org/10.3897/zookeys.929.35984

Cusi, J. C., Gagliardi-Urrutia, G., Brcko, I. C., Wake, D. B., & May, R. V. (2020). Taxonomic status of the Neotropical salamanders *Bolitoglossa altamazonica* and *Bolitoglossa peruviana* (Amphibia: Caudata: Plethodontidae), with the description of a new species from Northern Peru. Zootaxa, 4834(3), 365–406. https://doi.org/10.11646/zootaxa.4834.3.3

De Carvalho, T. R., & Giaretta, A. A. (2013). Taxonomic circumscription of *Adenomera martinezi* (Bokermann, 1956) (Anura: Leptodactylidae: Leptodactylinae) with the recognition of a new cryptic taxon through a bioacoustic approach. Zootaxa, 3701(2), 207. https://doi.org/10.11646/zootaxa.3701.2.5

Elmer, K. R., & Cannatella, D. C. (2008). Three new species of leaflitter frogs from the upper Amazon forests: Cryptic diversity within *Pristimantis* "*ockendeni*" (Anura: Strabomantidae) in Ecuador. Zootaxa, 1784(1), 11. https://doi.org/10.11646/zootaxa.1784.1.2

Firkowski, C. R., Bornschein, M. R., Ribeiro, L. F., & Pie, M. R. (2016). Species delimitation, phylogeny and evolutionary demography of co-distributed, montane frogs in the southern Brazilian Atlantic Forest. Molecular Phylogenetics and Evolution, 100, 345–360. https://doi.org/10.1016/j.ympev.2016.04.023

Flores Flores, E. (2018). Ampliación del rango de distribución y nuevos antecedentes del hábitat de dos especies de ranas del género *Alsodes* Bell, 1843 (Amphibia, Anura, Alsodidae) en la Cordillera de Nahuelbuta, sur de Chile. Biodiversity and Natural History (4), 1, 1-7.

Fouquet, A., Santana Cassini, C., Fernando Baptista Haddad, C., Pech, N., & Trefaut Rodrigues, M. (2014). Species delimitation, patterns of diversification and historical biogeography of the Neotropical frog genus *Adenomera* (Anura, Leptodactylidae). Journal of Biogeography, 41(5), 855–870. https://doi.org/10.1111/jbi.12250

Gehara, M., Canedo, C., Haddad, C. F. B., & Vences, M. (2013). From widespread to microendemic: Molecular and acoustic analyses show that *Ischnocnema guentheri* (Amphibia: Brachycephalidae) is endemic to Rio de Janeiro, Brazil. Conservation Genetics, 14(5), 973–982. https://doi.org/10.1007/s10592-013-0488-5

González, V. R., Schettino, L. R., Mancina, C. A., & Iturriaga, M. (2014). Amphibians Of Cuba: Checklist and geographic distributions. Smithsonian Herpetological Information Service, 145, 1–48. https://doi.org/10.5479/si.23317515.145.1

Higuera -Rojas, D. F., & Carvajal-Cogollo, J. E. (2021). Diet of Dendropsophus molitor (Anura: Hylidae) in a High-Andean agricultural ecosystem, Colombia. Universitas Scientiarum, 26(1), 119–137. https://doi.org/10.11144/Javeriana.SC26-1.dodm

Itgen, M. W., Sessions, S. K., Wilson, L. D., & Townsend, J. H. (2020). Integrative Systematic Revision of *Bolitoglossa celaque* (Caudata: Plethodontidae), with a new species from the Lenca Highlands of Honduras. Herpetological Monographs, 33(1), 48. https://doi.org/10.1655/HERPMONOGRAPHS-D-19-00001.1

Marin, C. M., Molina-Zuluaga, C., Restrepo, A., Cano, E., & Daza, J. M. (2018). A new species of *Leucostethus* (Anura: Dendrobatidae) from the eastern versant of the Central Cordillera of Colombia and the phylogenetic status of Colostethus fraterdanieli. Zootaxa, 4461(3), 359. https://doi.org/10.11646/zootaxa.4464.3.3

Mendelson, J. R., Mulcahy, D. G., Williams, T. S., & Sites Jr., J. W. (2011). A phylogeny and evolutionary natural history of mesoamerican toads (Anura: Bufonidae: *Incilius*) based on morphology, life history, and molecular data. Zootaxa, 3138(1), 1. https://doi.org/10.11646/zootaxa.3138.1.1

Meza-Joya, F. L., Hernández-Jaimes, C., & Ramos-Pallares, E. (2017). A new species of Salamander (Caudata, Plethodontidae, *Bolitoglossa*) from Serranía de los Yariguíes, Colombia. Zootaxa, 4294(1), 93. https://doi.org/10.11646/zootaxa.4294.1.4

Mulcahy, D. G., & Mendelson, J. R. (2000). Phylogeography and Speciation of the Morphologically Variable, Widespread Species *Bufo valliceps*, Based on Molecular Evidence from mtDNA. Molecular Phylogenetics and Evolution, 17(2), 173–189. https://doi.org/10.1006/mpev.2000.0827

Mulcahy, D. G., Morrill, B. H., & Mendelson, J. R. (2006). Historical biogeography of lowland species of toads (*Bufo*) across the Trans-Mexican Neovolcanic Belt and the Isthmus of Tehuantepec. Journal of Biogeography, 33(11), 1889–1904. https://doi.org/10.1111/j.1365-2699.2006.01546.x

Nuñez, J. J., Rabanal, F. E., & Formas, J. R. (2012). Description of a new species of *Eupsophus* (Amphibia: Neobatrachia) from the Valdivian Coastal range, Southern Chile: an integrative taxonomic approach. Zootaxa, 3305(1), 53. https://doi.org/10.11646/zootaxa.3305.1.3

Peters, J. A. (1973). The frog genus *Atelopus* in Ecuador (Anura: Bufonidae). Smithsonian Contributions to Zoology, 145, 1–49. https://doi.org/10.5479/si.00810282.145

Ramírez, J., Jaramillo, C., Lindquist, E., Crawford, A., & Ibáñez, R. (2020). Recent and Rapid Radiation of the Highly Endangered Harlequin Frogs (Atelopus) into Central America Inferred from Mitochondrial DNA Sequences. *Diversity*, *12*(9), 360. https://doi.org/10.3390/d12090360

Rebouças, R., & Solé, M. (2015). Diet of *Adenomera thomei* (Almeida and Angulo, 2006) (Anura: Leptodactylidae) from a rubber tree plantation in southern Bahia, Brazil. Studies on Neotropical Fauna and Environment, 50(2), 73–79. https://doi.org/10.1080/01650521.2015.1048613

Reis, S. F., Clemente-Carvalho, R. B. G., Santos, C. M. S. F. F., Lopes, R. T., Von Zuben, F. J., Laborda, P. R., & Perez, S. I. (2021). Skull diversity and evolution in miniaturized amphibians, genus *Brachycephalus* (Anura: Brachycephalidae). The Anatomical Record, 304(6), 1329–1343. https://doi.org/10.1002/ar.24554

Sanches, P. R., Pedroso-Santos, F., & Costa-Campos, C. E. (2019). Diet of *Adenomera hylaedactyla* (Cope, 1868) (Anura: Leptodactylidae) from an urban area in southern Amapá, eastern Amazon. Herpetology Notes,12: 841-845.

Taucce, P. P. G., Canedo, C., & Haddad, C. F. B. (2018). Two New Species of *Ischnocnema* (Anura: Brachycephalidae) from Southeastern Brazil and their Phylogenetic Position within the *I. guentheri* Series. Herpetological Monographs, 32(1), 1–21. https://doi.org/10.1655/HERPMONOGRAPHS-D-16-00014.1

Vigle, G. O., Coloma, L. A., Santos, J. C., Hernandez-Nieto, S., Ortega-Andrade, H. M., Paluh, D. J., & Read, M. (2020). A new species of *Leucostethus* (Anura: Dendrobatidae) from the Cordillera Mache-Chindul in northwestern Ecuador, with comments on similar Colostethus and Hyloxalus. *Zootaxa*, *4896*(3). https://doi.org/10.11646/zootaxa.4896.3.2

**Species not included in the phylogenetic analyses.**

*Allobates_tinae*
*Brachycephalus_crispus*
*Brachycephalus_sulfuratus*
*Melanophryniscus_biancae*
*Melanophryniscus_milanoi*
*Melanophryniscus_xanthostomus*
*Rhinella_horribilis*
*Andinobates_victimatus*
*Leucostethus_jota*
*Adenomera_chicomendesi*
*Pristimantis_ardilae*
*Bolitoglossa_awajun*
*Bolitoglossa_copinhorum*
*Bolitoglossa_yariguiensis*
*Thorius_maxillabrochus*

**Supplementary figures**

**Figure S1.1.** Number of observations/sequences per grid cell (a,c), and amphibian genetic diversity patterns in the Neotropics (b,d). The map uses equal-area grid cells of 150 (a,b) and 250 km (c,d).

**Figure S1.2.** (a) Simulate evolution along each branch of the tree following Brownian Motion model. A Brownian Motion model assumes that traits evolve as a random walk along the length of a branch in a phylogeny. (b) Simulating Brownian Motion on our tree with fastBM() function in phytools.

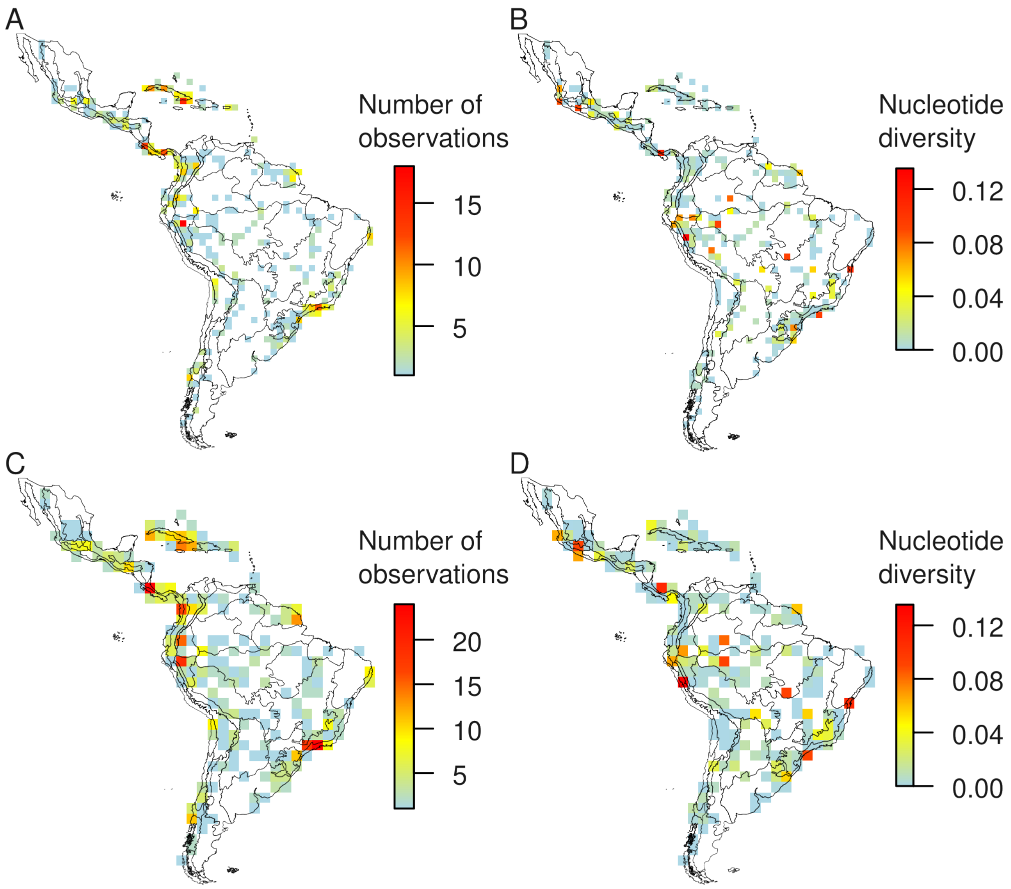


**Figure S1.1.** Number of observations/sequences per grid cell (a,c), and amphibian genetic diversity patterns in the Neotropics (b,d). The map uses equal-area grid cells of 150 (a,b) and 250 km (c,d).

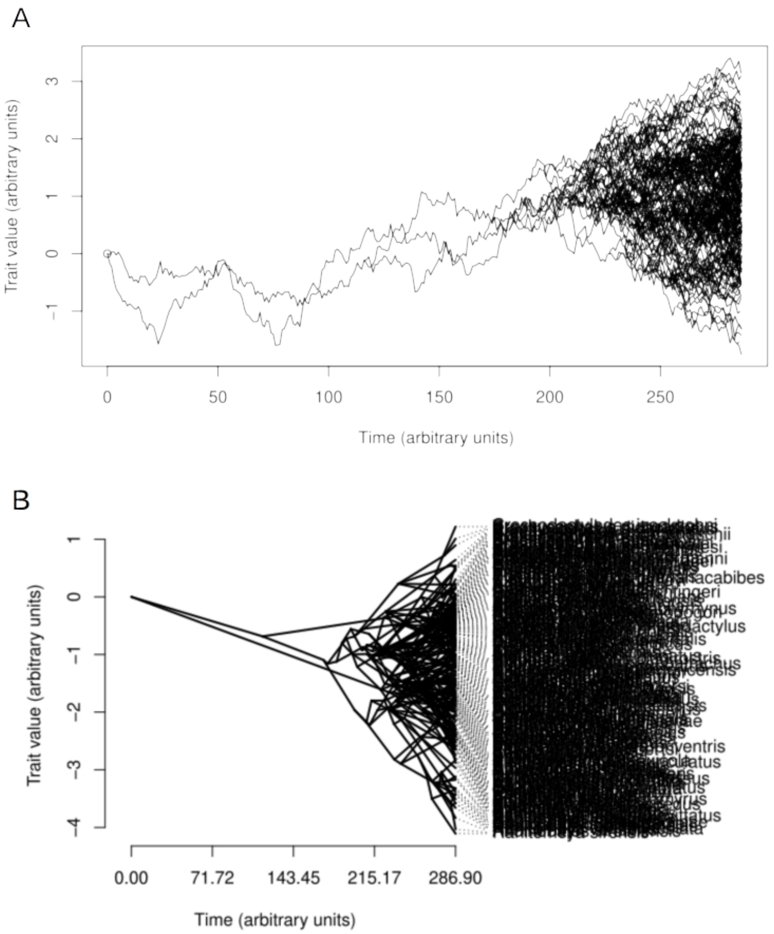


**Figure S1.2.** (a) Simulate evolution along each branch of the tree following Brownian Motion model. A Brownian Motion model assumes that traits evolve as a random walk along the length of a branch in a phylogeny. (b) Simulating Brownian Motion on our tree with fastBM() function in *phytools*.

**Appendix S2** Supplementary results

### Supplementary tables

**Table S2.1.** Results of Phylogenetic Linear (pi) and Logistic (IBD – in blue) regression models using *phylolm* R package. * This table can be found in the figshare online repository https://doi.org/10.6084/m9.figshare.23520384.v1

### Supplementary figures

**Figure S2.1.** Basic histogram showing the number of species associated with the number of CytB sequences.

**Figure S2.2.** Map of occurrences associated with mtDNA sequences from Neotropical frogs and salamanders.

**Figure S2.3.** Isolation-by-Distance (IBD) patterns of amphibians by Neotropical region.

**Figure S2.4.** Random forest results for Anura and Caudata separately show the top predictors of nucleotide diversity, percentage of variance explained of the models and the relative importance is also shown.

**Figure S2.5.** Random forest results for Anura and Caudata separately showing the top predictors of genetic structure. Out of bag (OOB) estimates percentage of error are also shown.

**Figure S2.6.** Random forest results for all amphibians using reduced datasets showing the top predictors of nucleotide diversity, A) at least 10 sequences per species, B) at least 400 bp in each species alignment, and C) both at least 10 sequences per species and 400 bp in each species alignment. The percentage of variance explained, and the relative importance is also shown for each model.

**Figure S2.7.** Random forest results for all amphibians using reduced datasets showing the top predictors of genetic structure, A) at least 10 sequences per species, B) at least 400 bp in each species alignment, and C) both at least 10 sequences per species and 400 bp in each species alignment. Out of bag (OOB) estimates percentage of error are also shown.

**Figure S2.8.** Phylogeny of Neotropical frogs and salamanders, with nucleotide diversity ($\pi$) traced across branches. The phylogeny is based on CytB DNA sequences data for 241 species. Nucleotide diversity box plots per taxonomic family are indicated to the right.

**Figure S2.9.** Phylogenetic signal for nucleotide diversity ($\pi$) and population structure (IBD). A) Likelihood surface of the phylogenetic signal based on Pagel's lambda ($\lambda$) for $\pi$. A $\lambda$ value = 0.746 indicates a correlation between species close to a Brownian motion model of evolution.
B) Phylogenetic signal D for population structure (binary variable). A value close to 1 (D = 0.763) indicates that IBD is randomly shuffled relative to the tips of the phylogeny (phylogenetic randomness).

**Figure S2.10.** Patterns in the distribution of genetic diversity ($\pi$) of Neotropical amphibians over the geographical range size in logarithmic scale for A) Anura (salmon) and Caudata (turquoise) combined, B) Anura, and C) Caudata. The equation of each model is shown on the plots.

**Figure S2.11.** Minimum elevational patterns of genetic diversity ($\pi$) in Neotropical amphibians for A) Anura (salmon) and Caudata (turquoise) combined, B) Anura, and C) Caudata. The equation of each model is shown on the plots.

**Figure S2.12.** Latitude as predictor of genetic structure or Isolation-by-Distance (IBD) in Neotropical amphibians. A) Mean Latitude, B) Maxium Latitude. No IBD (red), IBD (green). Each box-whisker plot indicates the median (bold lines), and the interquartile range (boxes).

**Figure S2.13.** Isolation-by-Distance (IBD) patterns of Neotropical amphibians by activity.

**Figure S2.14.** Body size as predictor of genetic structure or Isolation-by-Distance (IBD) in Caudata (A) and Anura (B). No IBD (red), IBD (green). Each box-whisker plot indicates the median (bold lines), the interquartile range (boxes), white dots represent outliers.

**Figure S2.15.** Mean (A-B) and minimum elevation (C-D) as predictors of genetic structure or Isolation-by-distance (IBD), for frogs (A, C) and salamanders (B, D) respectively. Each box-whisker plot indicates the median (bold lines), the interquartile range (boxes), white dots represent outliers.

**Figure S2.16.** Posterior distribution of Isolation-by-Distance (IBD) across all of predictors, plotting their estimate and associated 95% credible intervals (CIs). The 95% CIs do not cross zero for semi-aquatic microhabitat and Antillean and Mesoamerica biogeographic units, suggesting that these predictors are statistically significant.

**Figure S2.17.** Trace and density of the intercept and predictors of nucleotide diversity. Trace plots as diagnostics that allow judge whether the MCMC chains in the MCMCglmm analyses are likely to have converged on stationary distributions. The traces indicate that chains have mixed well and converged.

**Figure S2.18.** Trace and density of the intercept and predictors of population structure (IBD, second best model). Trace plots as diagnostics that allow judge whether the MCMC chains in the MCMCglmm analyses are likely to have converged on stationary distributions. The traces indicate that chains have mixed well and converged.

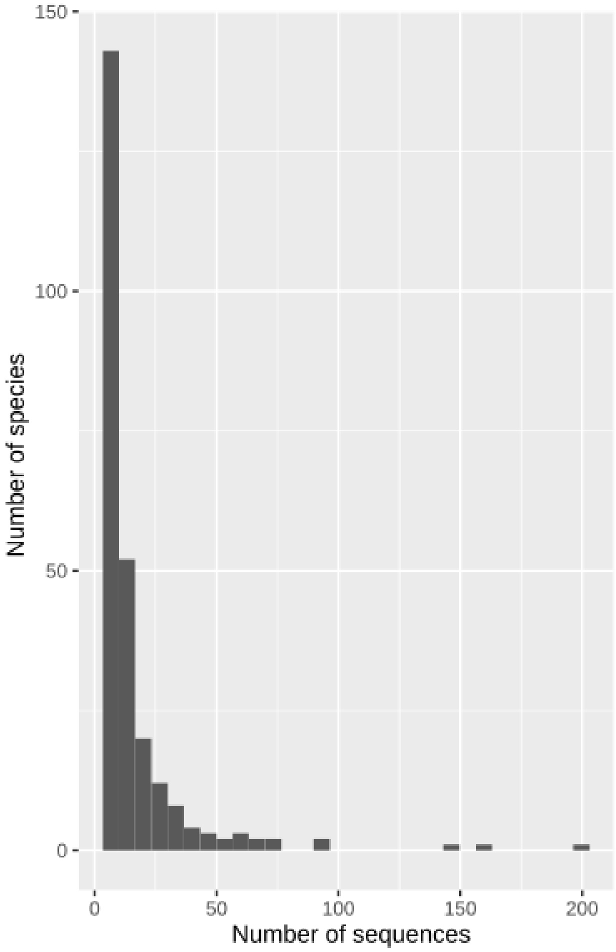


**Figure S2.1.** Basic histogram showing the number of species associated with the number of CytB sequences.

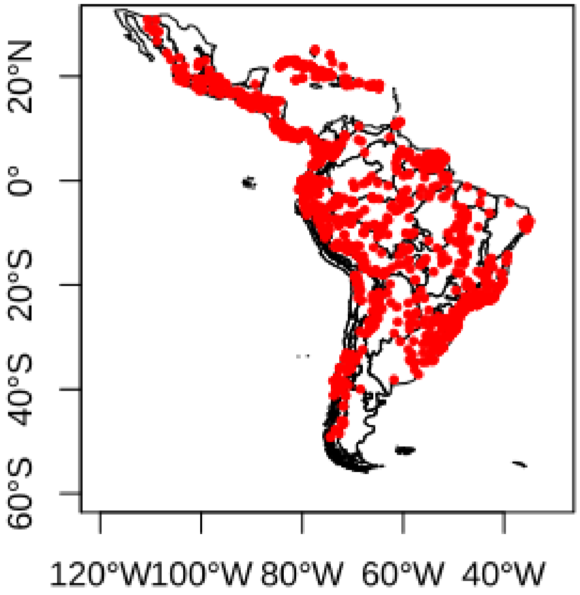


**Figure S2.2.** Map of occurrences associated with mtDNA sequences from Neotropical frogs and salamanders.

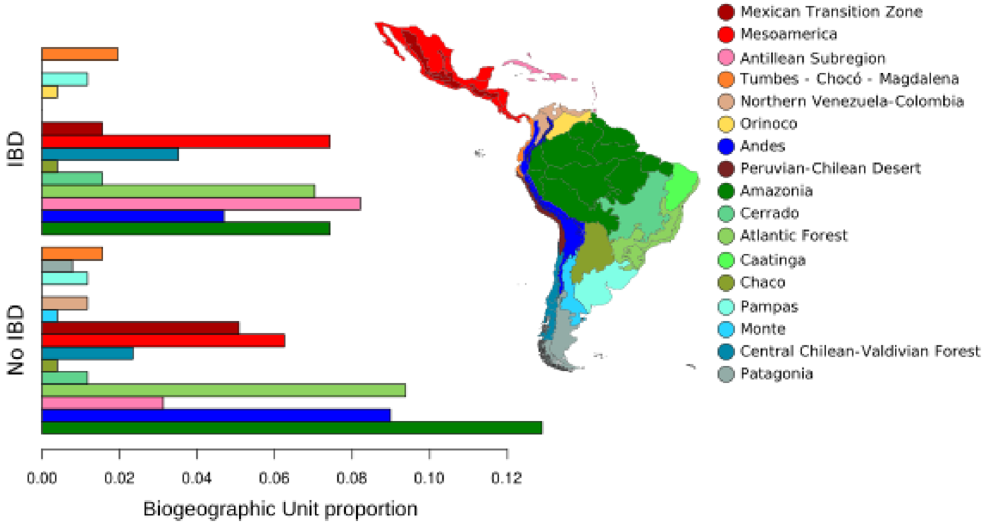


**Figure S2.3.** Isolation-by-Distance (IBD) patterns of amphibians by Neotropical region.

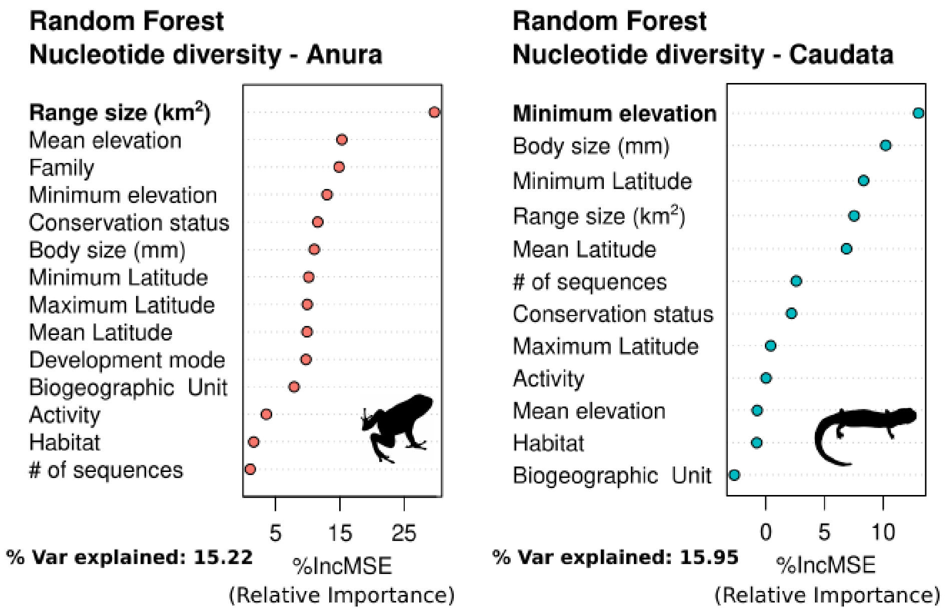


**Figure S2.4.** Random forest results for Anura and Caudata separately show the top predictors of nucleotide diversity, percentage of variance explained of the models and the relative importance is also shown.

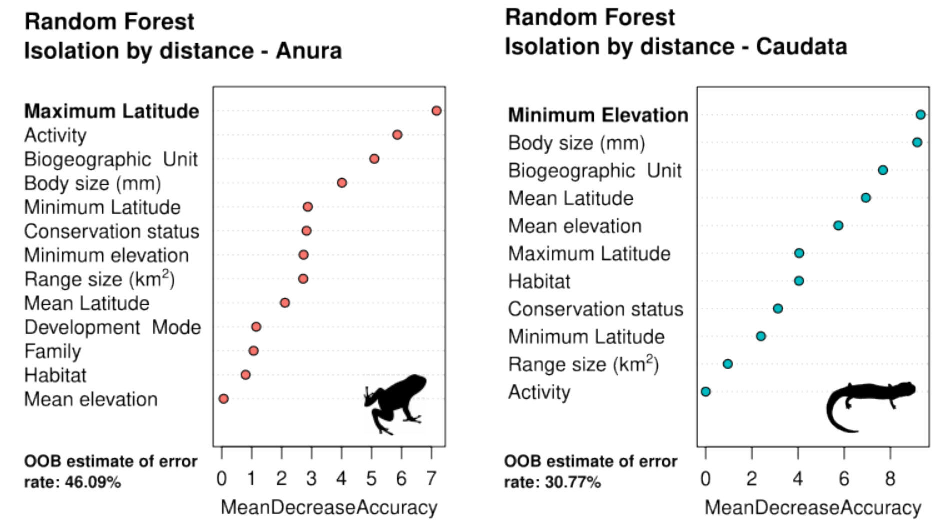


**Figure S2.5.** Random forest results for Anura and Caudata separately showing the top predictors of genetic structure. Out of bag (OOB) estimates percentage of error are also shown.

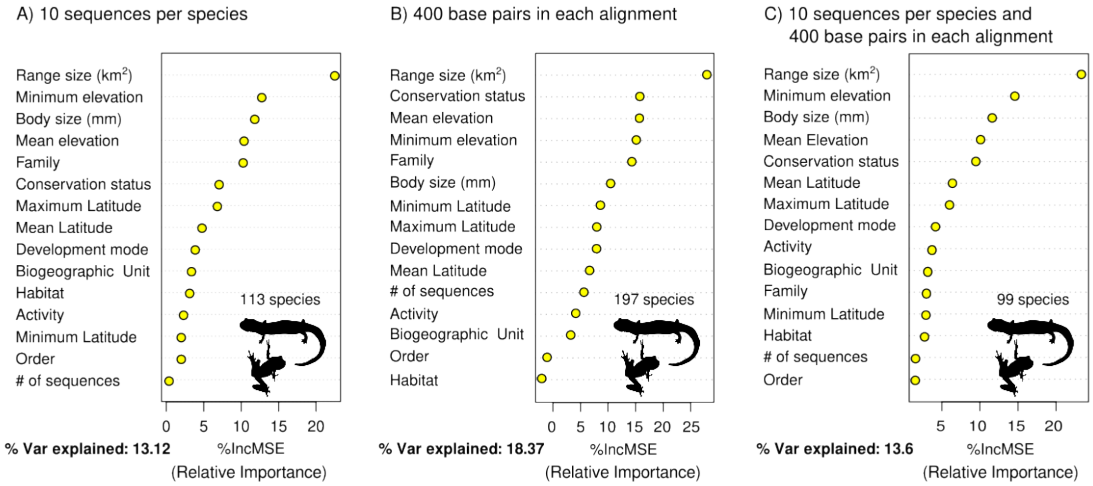


**Figure S2.6.** Random forest results for all amphibians using reduced datasets showing the top predictors of nucleotide diversity, A) at least 10 sequences per species, B) at least 400 bp in each species alignment, and C) both at least 10 sequences per species and 400 bp in each species alignment. The percentage of variance explained, and the relative importance is also shown for each model.

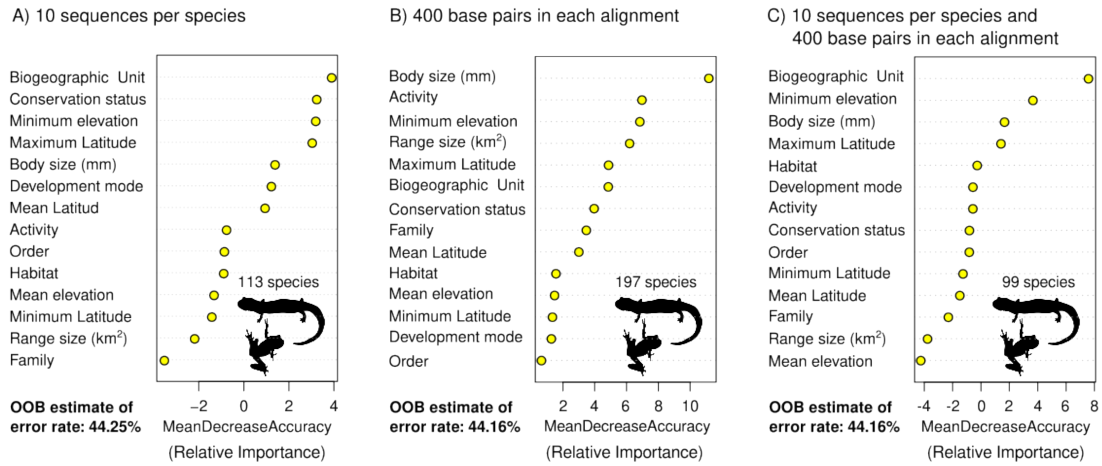


**Figure S2.7.** Random forest results for all amphibians using reduced datasets showing the top predictors of genetic structure, A) at least 10 sequences per species, B) at least 400 bp in each species alignment, and C) both at least 10 sequences per species and 400 bp in each species alignment. Out of bag (OOB) estimates percentage of error are also shown.

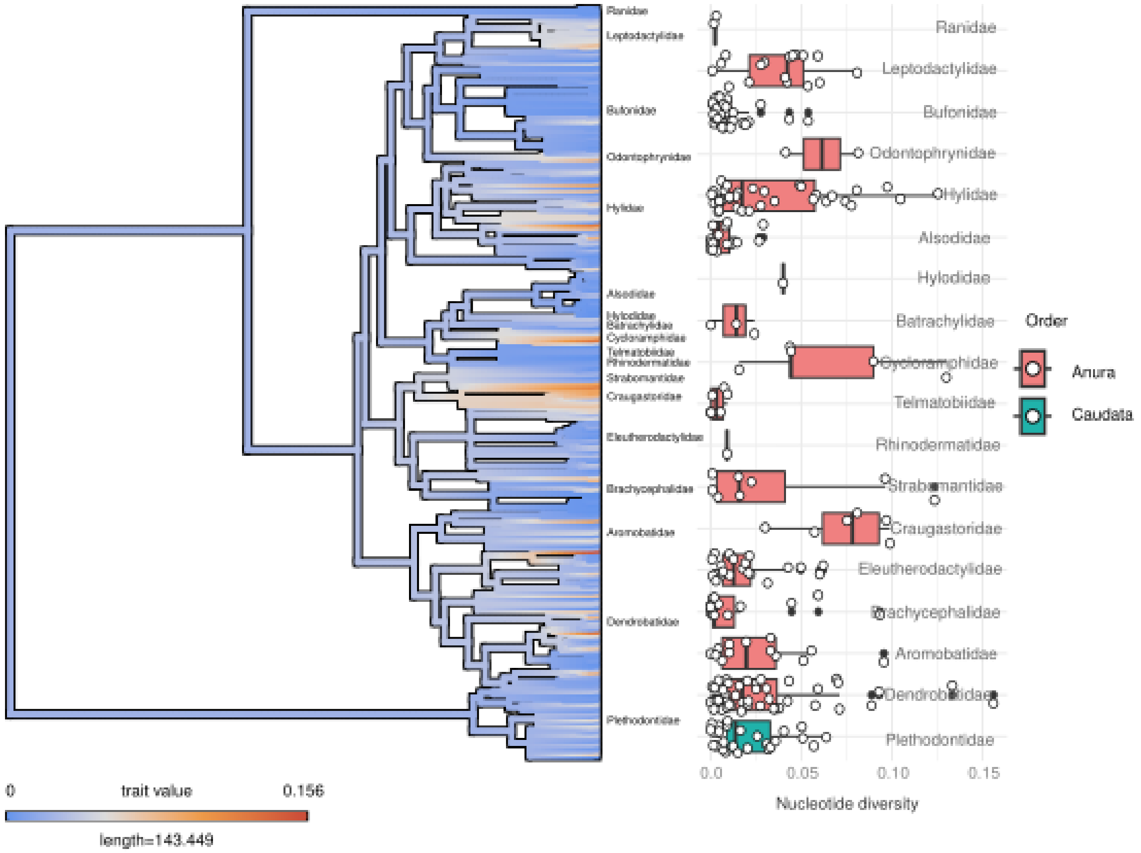


**Figure S2.8.** Phylogeny of Neotropical frogs and salamanders, with nucleotide diversity ($\pi$) traced across branches. The phylogeny is based on CytB DNA sequences data for 241 species. Nucleotide diversity box plots per taxonomic family are indicated to the right.

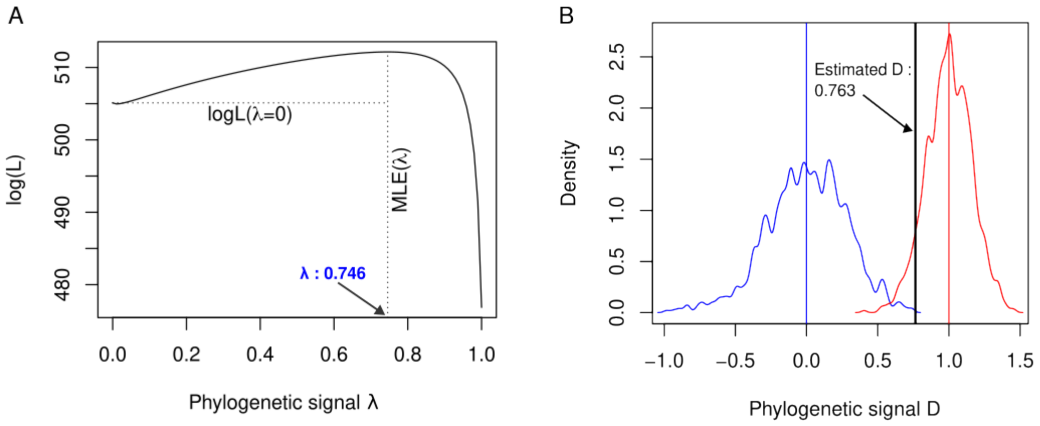


**Figure S2.9.** Phylogenetic signal for nucleotide diversity ($\pi$) and population structure (IBD). A) Likelihood surface of the phylogenetic signal based on Pagel's lambda ($\lambda$) for $\pi$. A $\lambda$ value = 0.746 indicates a correlation between species close to a Brownian motion model of evolution. B) Phylogenetic signal D for population structure (binary variable). A value close to 1 (D = 0.763) indicates that IBD is randomly shuffled relative to the tips of the phylogeny (phylogenetic randomness).

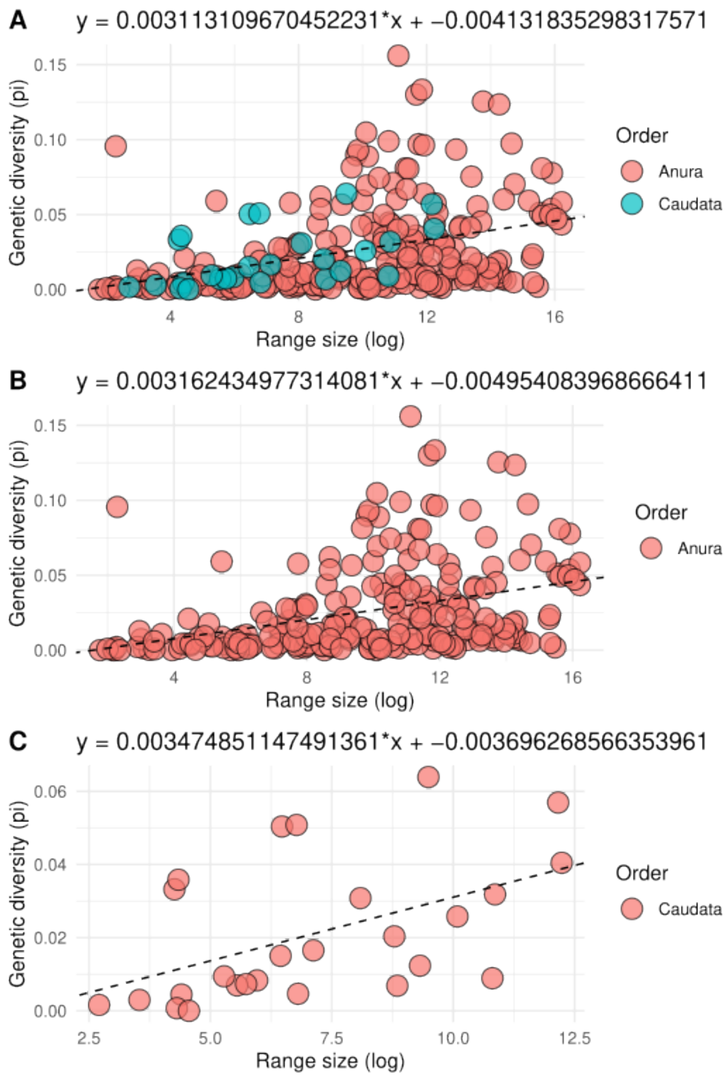


**Figure S2.10.** Patterns in the distribution of genetic diversity ($\pi$) of Neotropical amphibians over the geographical range size in logarithmic scale for A) Anura (salmon) and Caudata (turquoise) combined, B) Anura, and C) Caudata. The equation of each model is shown on the plots.

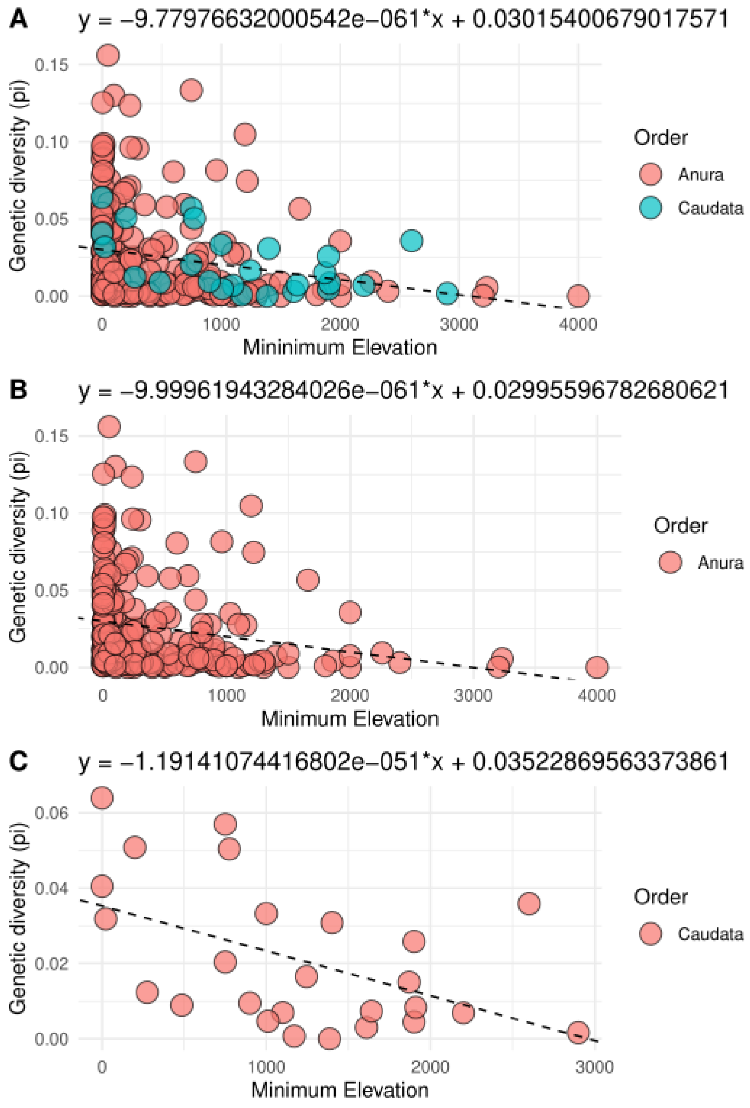


**Figure S2.11.** Minimum elevation patterns of genetic diversity ($\pi$) in Neotropical amphibians for A) Anura (salmon) and Caudata (turquoise) combined, B) Anura, and C) Caudata. The equation of each model is shown on the plots.

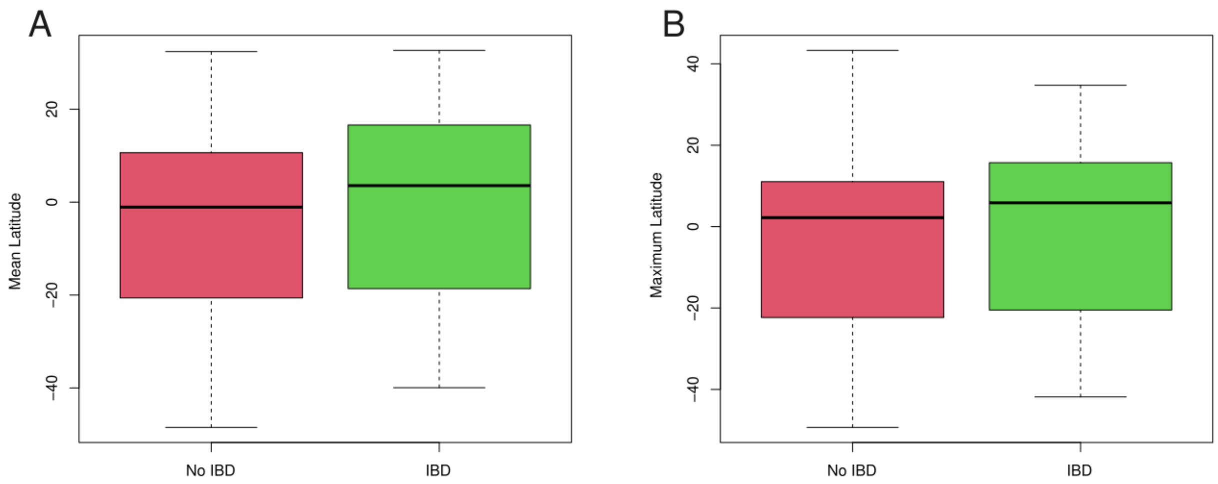


**Figure S2.12.** Latitude as predictor of genetic structure or Isolation-by-Distance (IBD) in Neotropical amphibians. A) Mean Latitude, B) Maxium Latitude. No IBD (red), IBD (green). Each box-whisker plot indicates the median (bold lines), and the interquartile range (boxes).

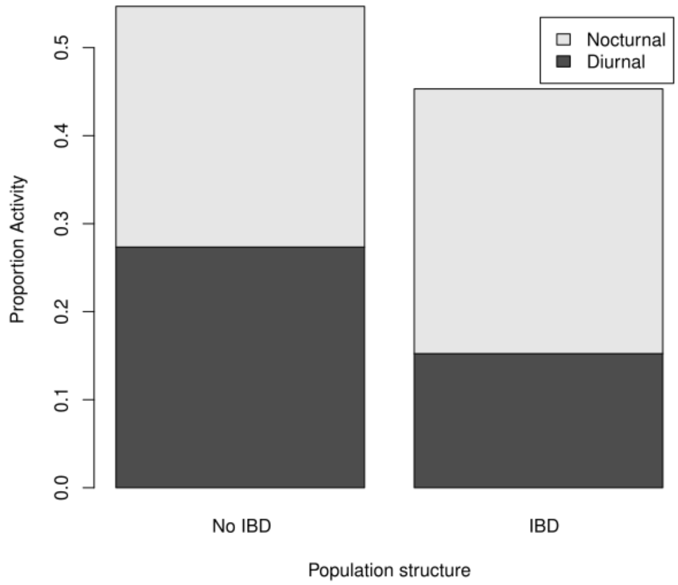


**Figure S2.13.** Isolation-by-Distance (IBD) patterns of Neotropical amphibians by activity.

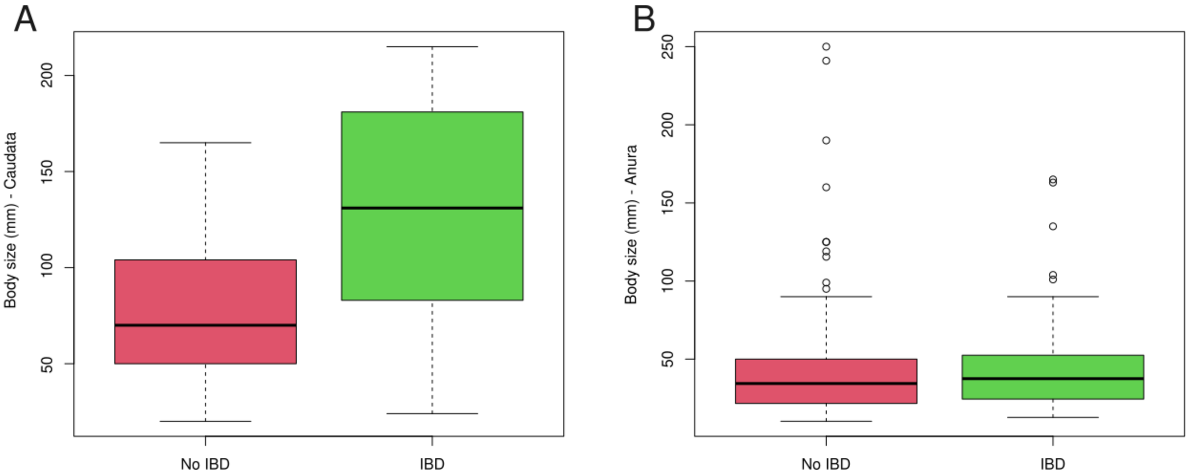


**Figure S2.14.** Body size as predictor of genetic structure or Isolation-by-Distance (IBD) in Caudata (A) and Anura (B). No IBD (red), IBD (green). Each box-whisker plot indicates the median (bold lines), the interquartile range (boxes), white dots represent outliers.

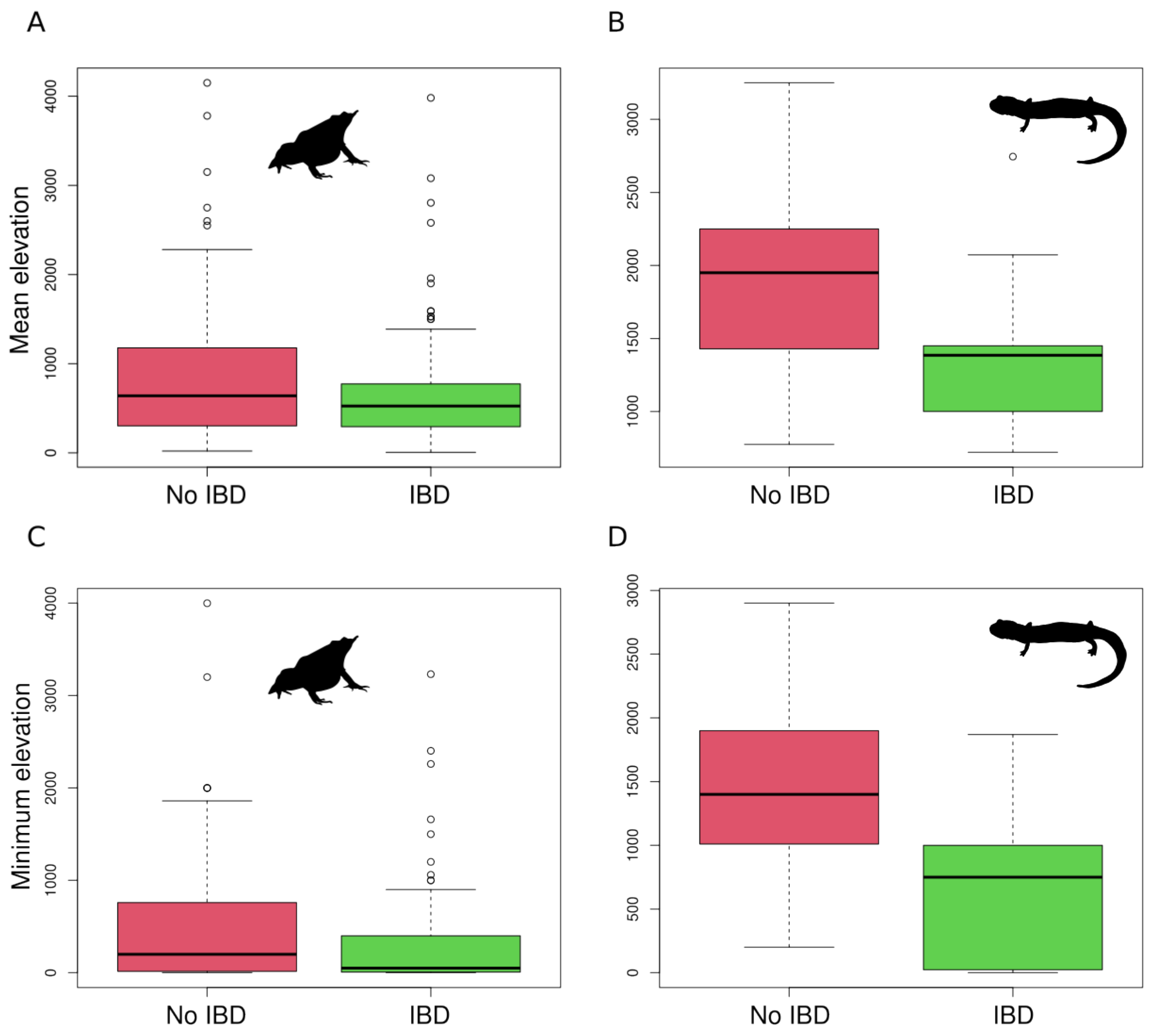


**Figure S2.15.** Mean (A-B) and minimum elevation (C-D) as predictors of genetic structure or Isolation-by-distance (IBD), for frogs (A, C) and salamanders (B, D) respectively. Each box-whisker plot indicates the median (bold lines), the interquartile range (boxes), white dots represent outliers.

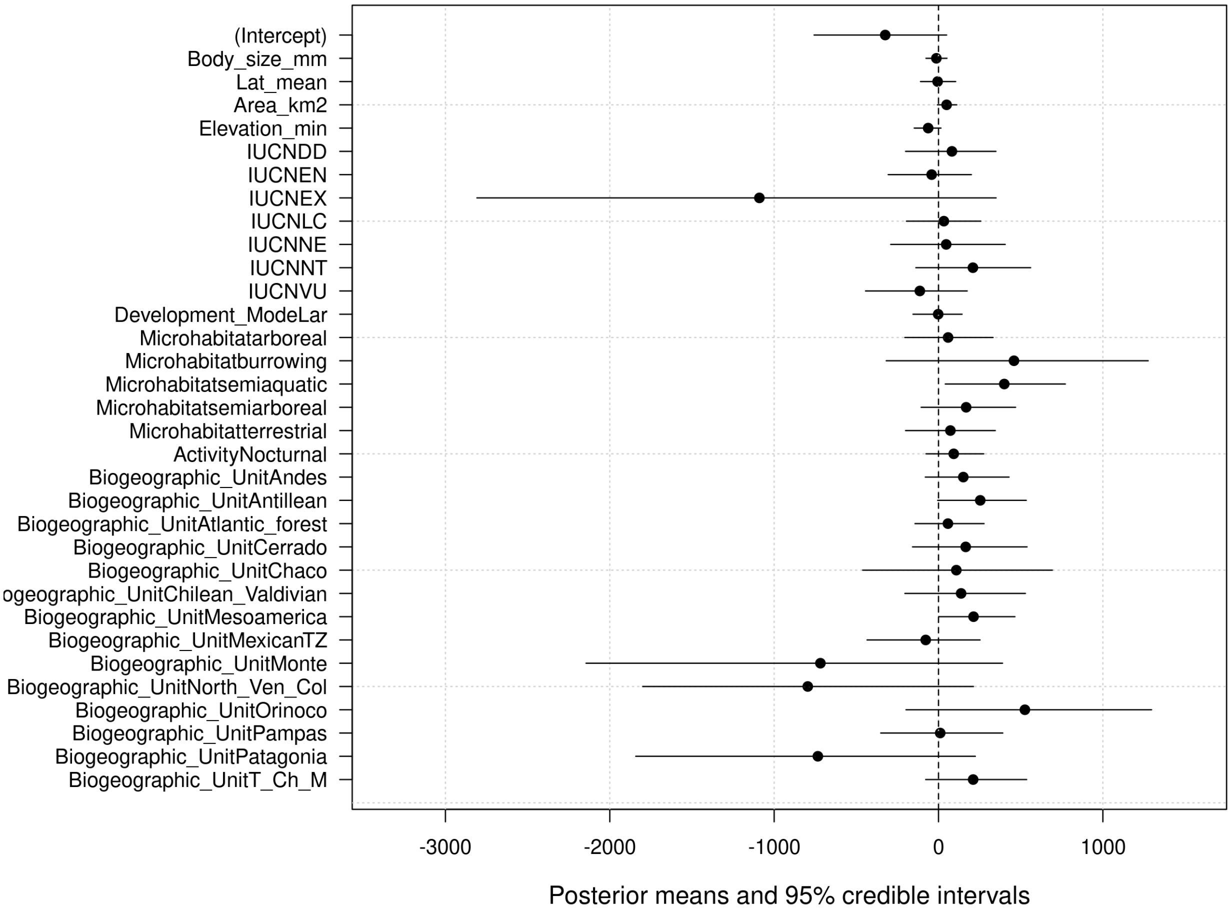


**Figure S2.16.** Posterior distribution of Isolation-by-Distance (IBD) across all of predictors, plotting their estimate and associated 95% credible intervals (CIs). The 95% CIs do not cross zero for semi-aquatic microhabitat and Antillean and Mesoamerica biogeographic units, suggesting that these predictors are statistically significant.

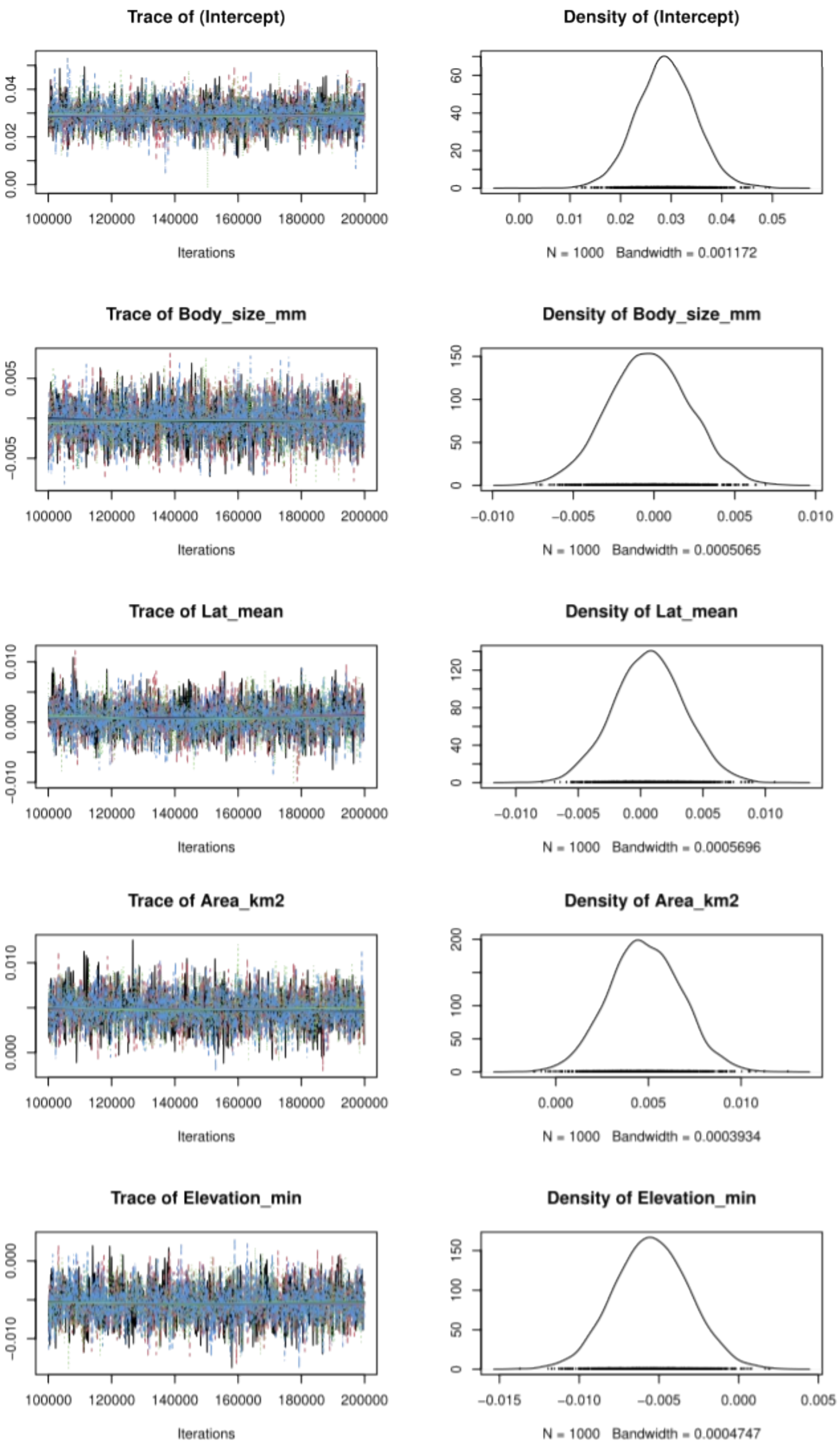


**Figure S2.17.** Trace and density of the intercept and predictors of nucleotide diversity. Trace plots as diagnostics that allow judge whether the MCMC chains in the MCMCglmm analyses are likely to have converged on stationary distributions. The traces indicate that chains have mixed well and converged.

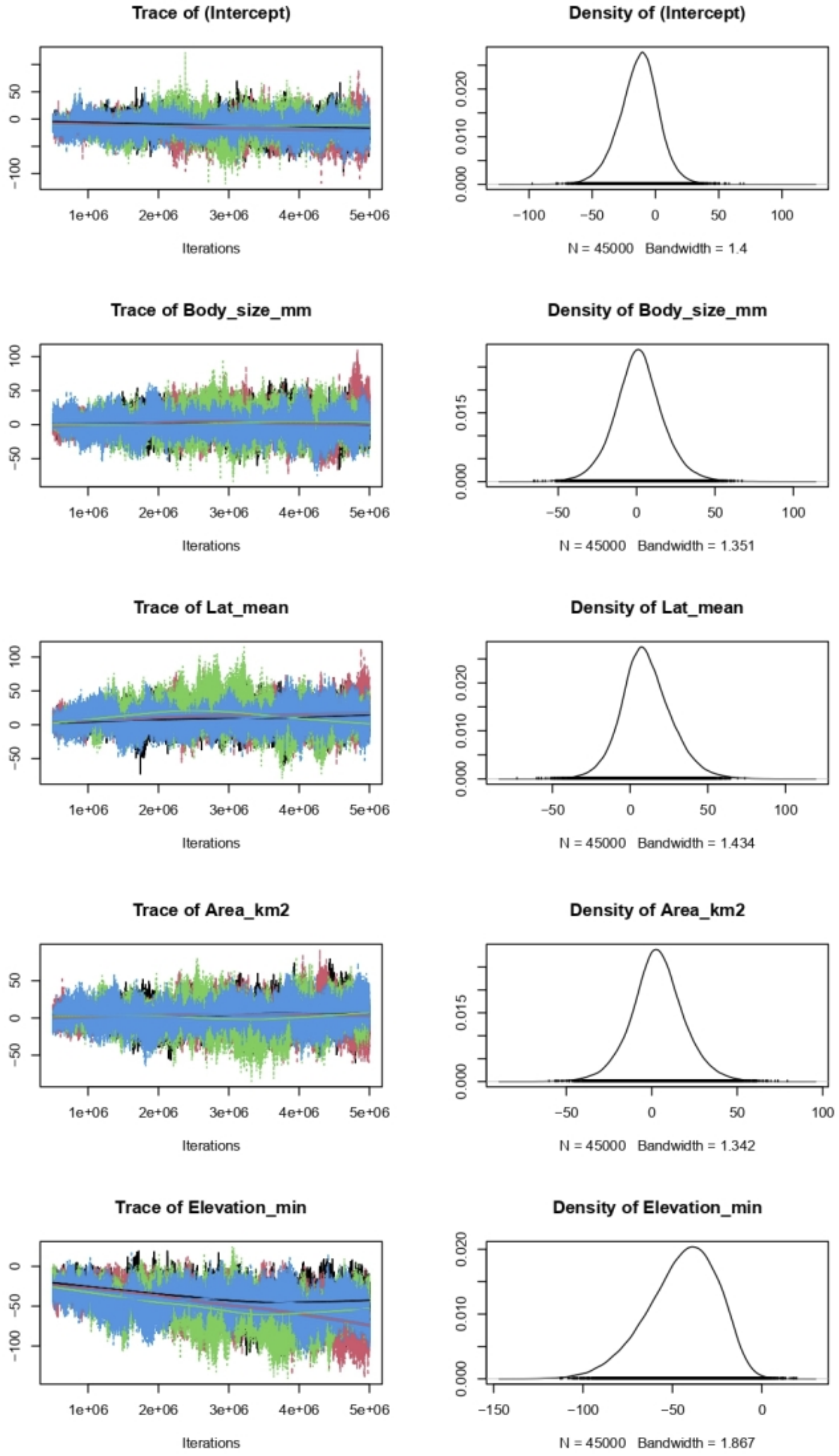


**Figure S2.18.** Trace and density of the intercept and predictors of population structure (IBD, second best model). Trace plots as diagnostics that allow judge whether the MCMC chains in the MCMCglmm analyses are likely to have converged on stationary distributions. The traces indicate that chains have mixed well and converged.

**# Data with no outliers ($\pi$)**

**Species outliers:**

*Cycloramphus eleutherodactylus* 0.13003375

*Ranitomeya sirensis* 0.13333333

*Silverstoneia nubicola* 0.15601375

*Boana balzani* 0.10472441

*Dendropsophus decipiens* 0.12541403

*Pristimantis altamazonicus* 0.12352352

**Phylogenetic signal results**

**# Nucleotide diversity**
Phylogenetic signal lambda : 0.745562
logL(lambda) : 512.146
LR(lambda=0) : 14.0725
P-value (based on LR test) : 0.000175892

**# Population structure**
Calculation of D statistic for the phylogenetic structure of a binary variable
Number of permutations : 1000
Estimated D : 0.7632384
Probability of E(D) resulting from no (random) phylogenetic structure : 0.081
Probability of E(D) resulting from Brownian phylogenetic structure : 0.001

**# Body size**
All - lambda = 0.978714, P = 9.34458e-40
Anura - lambda = 0.92981, P = 1.84678e-33
Caudata - lambda = 0.999934, P = 0.108221

**# Number of Sequences**
All - lambda = 0.219472, P =0.0334779
Anura - lambda = 0.140334, P = 0.0807153
Caudata - lambda = 6.6107e-05, P = 1

**# Mean Elevation**
All - lambda = 0.949644, P = 1.88135e-29
Anura - lambda = 0.919331, P = 7.06112e-24
Caudata - lambda = 6.6107e-05, P = 1

**# Minimum Elevation**

All - lambda = 0.941357, P = 5.46502e-23
Anura - lambda = 0.89705, P = 8.00148e-21
Caudata - lambda = 6.6107e-05, P = 1

**# Mean Latitude**
All - lambda = 0.999934, P = 2.84733e-70
Anura - lambda = 0.999934, P = 7.63998e-62
Caudata - lambda = 0.999934, P = 0.0377557

**# Range Size**
All - lambda = 0.403947, P = 0.000407788
Anura - lambda = 0.285667, P = 0.00101213
Caudata - lambda = 6.6107e-05, P = 1

**Appendix S3** Discussion – supplementary information

**Table S3.1.** Species complex or cryptic species analyzed in this study.

| **Species** | **Family** | **$\pi$ value** | **IBD** | **Reference** |
|---|---|---|---|---|
| *Pristimantis altamazonicus* | Strabomantidae | 0.1235 | Yes | Ortega-Andrade et al., 2017 |
| *Dendropsophus decipiens* | Hylidae | 0.1254 | Yes | Oliveira et al., 2021 |
| *Scinax squalirostris* | Hylidae | 0.0974 | Yes | Faria et al., 2013 |
| *Ischnocnema guentheri* | Brachycephalidae | 0.0934 | Yes | Gehara et al., 2013 |
| *Craugastor occidentalis* | Craugastoridae | 0.0970 | Yes | Streicher et al., 2014 |
| *Anomaloglossus degranvillei* | Aromobatidae | 0.0956 | Yes | Vacher et al., 2017 |
| *Phyllobates lugubris* | Dendrobatidae | 0.9290 | No | Marquez et al., 2020 |
| *Ranitomeya sirensis* | Dendrobatidae | 0.1333 | No | Brown et al., 2011 |
| *Silverstoneia nubicola* | *Dendrobatidae* | 0.1560 | No | Grant and Myers, 2013 |

### Supplementary figures

**Figure S3.1.** Relationship between geographic range size in logarithmic scale and IUCN conservation categories of Neotropical amphibians for Anura (salmon) and Caudata (turquoise) combined. Each box-whisker plot indicates the median (bold lines), the interquartile range (boxes), white dots represent an observation of species, and black-white dots represent outliers. A single bold line with a white dot represents a single observation. DD= Data Deficient, LC=Least Concern, NT=Near Threatened, VU= Vulnerable, EN=Endangered, CR= Critically Endangered, EX= Extinct, NE= Not Evaluated.

**Figure S3.2.** Genetic diversity ($\pi$) of Neotropical amphibians across IUCN conservation categories for A) Anura (salmon) and Caudata (turquoise) combined, B) Anura, and C) Caudata. Each box-whisker plot indicates the median (bold lines), the interquartile range (boxes), white dots represent an observation of $\pi$, and black-white dots represent outliers. A single bold line with a white dot represents a single observation. DD= Data Deficient, LC=Least Concern, NT=Near Threatened, VU= Vulnerable, EN=Endangered, CR= Critically Endangered, EX= Extinct, NE= Not Evaluated.

**Figure S3.3.** Nucleotide diversity is high in latitudes near the equator for Neotropical amphibians following the latitudinal gradient of species richness.

**Figure S3.4.** Genetic diversity is positively correlated with body size in salamanders.

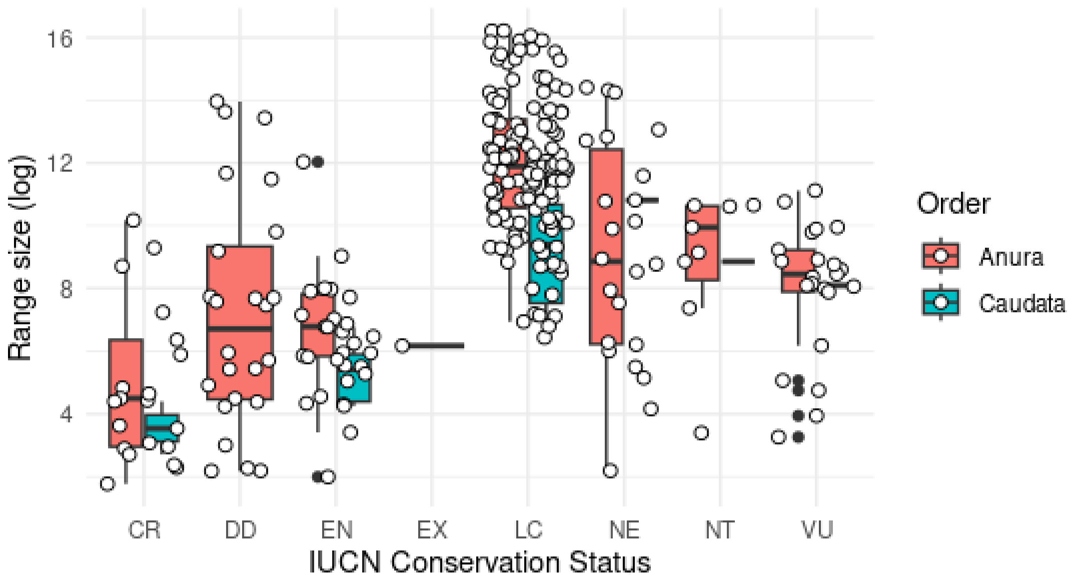


**Figure S3.1.** Relationship between geographic range size in logarithmic scale and IUCN conservation categories of Neotropical amphibians for Anura (salmon) and Caudata (turquoise) combined. Each box-whisker plot indicates the median (bold lines), the interquartile range (boxes), white dots represent an observation of species, and black-white dots represent outliers. A single bold line with a white dot represents a single observation. DD= Data Deficient, LC=Least Concern, NT=Near Threatened, VU= Vulnerable, EN=Endangered, CR= Critically Endangered, EX= Extinct, NE= Not Evaluated.

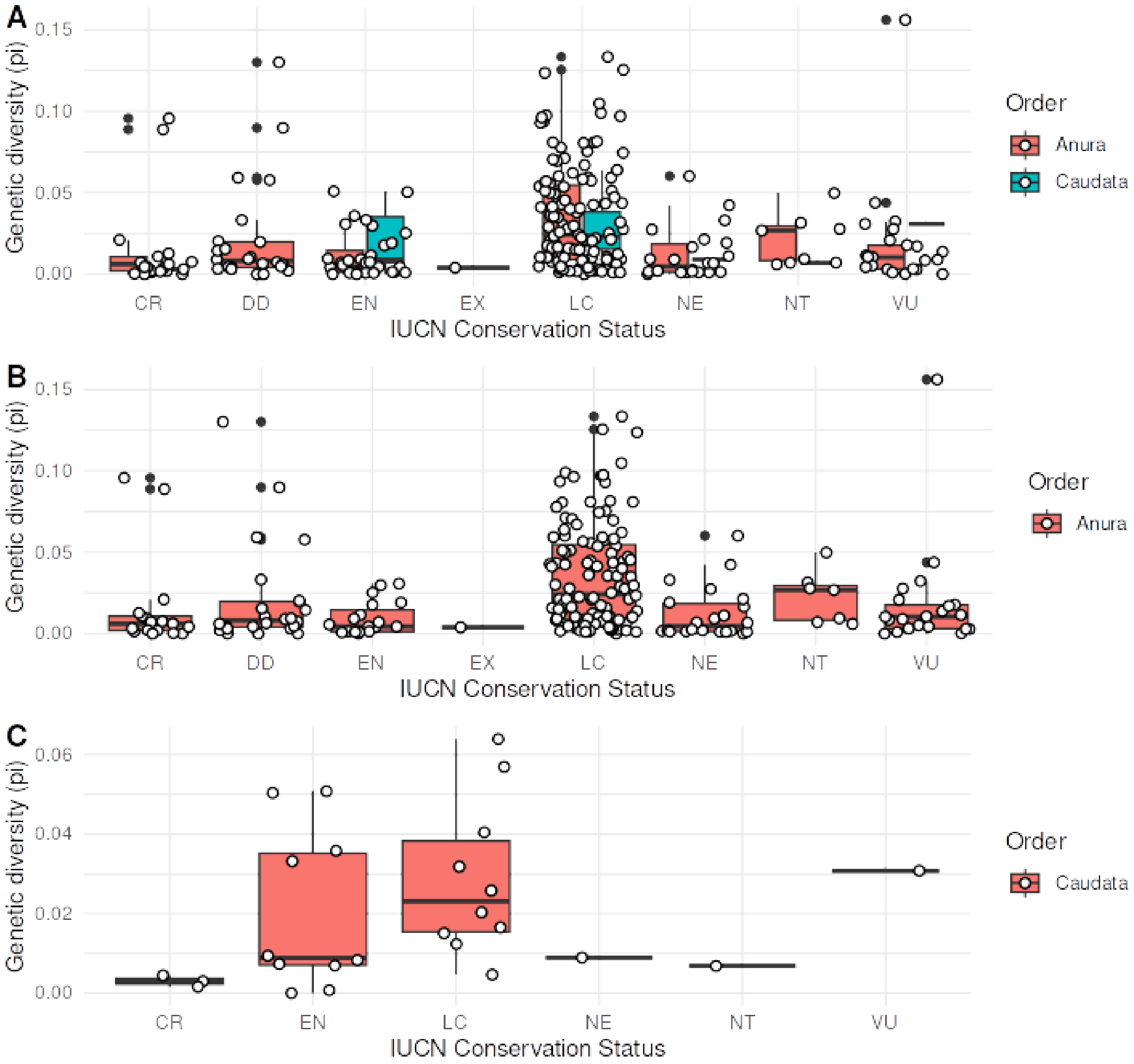


**Figure S3.2.** Genetic diversity ($\pi$) of Neotropical amphibians across IUCN conservation categories for A) Anura (salmon) and Caudata (turquoise) combined, B) Anura, and C) Caudata. Each box-whisker plot indicates the median (bold lines), the interquartile range (boxes), white dots represent an observation of $\pi$, and black-white dots represent outliers. A single bold line with a white dot represents a single observation. DD= Data Deficient, LC=Least Concern, NT=Near Threatened, VU= Vulnerable, EN=Endangered, CR= Critically Endangered, EX= Extinct, NE= Not Evaluated.

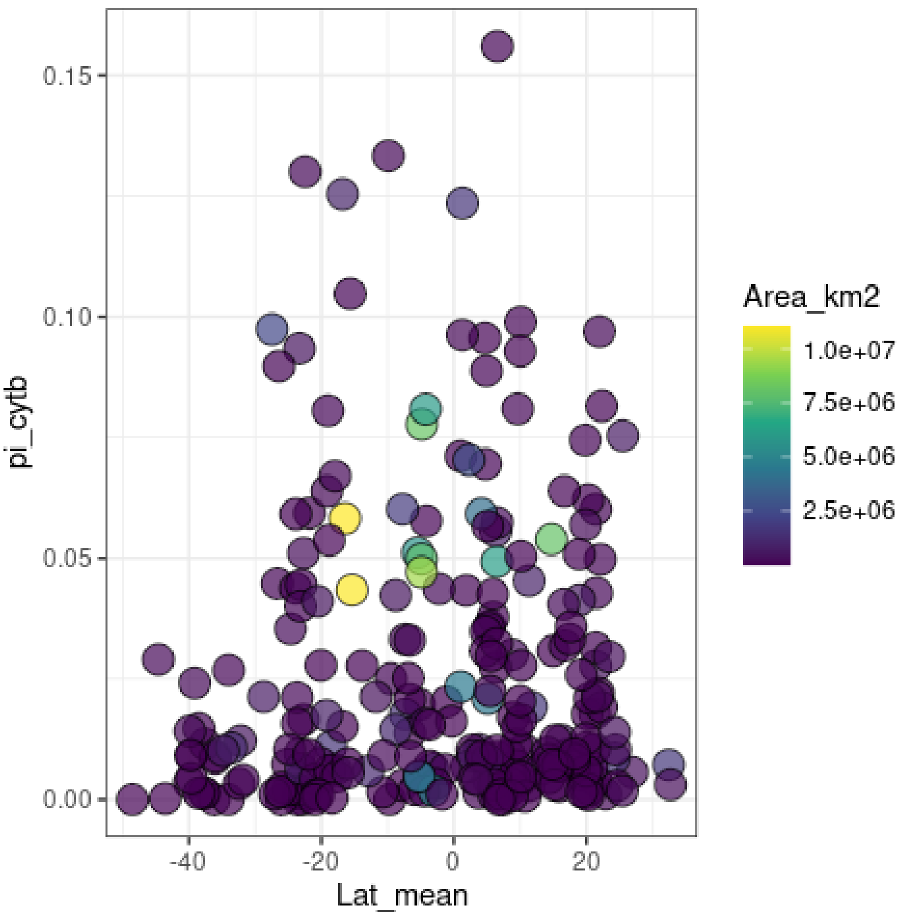


**Figure S3.3.** Nucleotide diversity is high in latitudes near the equator for Neotropical amphibians following the latitudinal gradient of species richness.

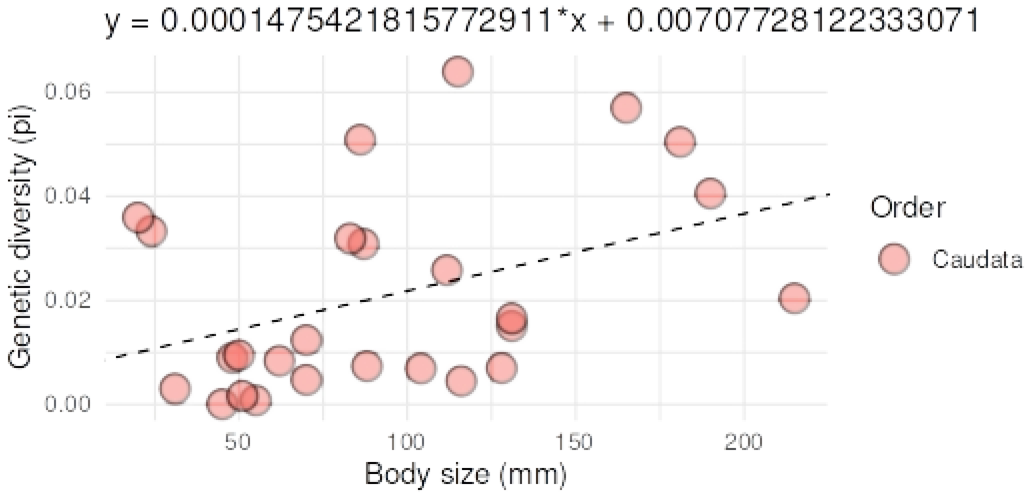


**Figure S3.4.** Genetic diversity is positively correlated with body size in salamanders.
